# An intrinsically disordered proline-rich domain acts in synergy with AGP-type glycosylation to control the protein fate of FLA4

**DOI:** 10.64898/2026.10.06.756988

**Authors:** Chaitra Hiremath, Hui Xue, Tuba Acet, Georg J. Seifert

**Affiliations:** Institute of Plant Biotechnology and Cell Biology, BOKU University, Vienna; Muthgasse 18 1190-Vienna, Austria; TA: Department of Occupational Health and Safety, Faculty of Health Sciences, Gümüşhane University, Gümüşhane, Türkiye

## Abstract

The functional roles of glycans attached to the proline (Pro) rich region of arabinogalactan proteins are largely elusive. We have tested the possibility of type II arabinogalactans acting as protein fate determinants for the FASCICLIN LIKE ARABINOGALACTAN PROTEIN 4 (FLA4). Pro-to-Ala substitutions in the carboxy-terminal region caused *in planta* protein destabilization in an *ECHIDNA*-dependent trans-Golgi compartment and plasma membrane shedding in an aspartate protease-dependent manner. The introduction of diverse glycomodules to the FLA4 carboxy-terminus counteracted the stability loss, with AGP type glycosylation motifs more efficient than *N-*glycosylation signals and extensin structures. The resistance of FLA4 stability and localization to genetic interference with *O-*glycosylation suggested that additional protein properties, such as intrinsic disorder and secondary structure were crucial for FLA4 stability. Consistently, disorder-maintaining site directed mutants maintained FLA4 stability in a glycosylation-independent manner. Our data support a role for proline-rich regions in determining protein fate by generating intrinsically disordered domains in the peptide chain and providing acceptor sites for *O*-glycosylation.

**Significance statement:** We tested the hypothetical role of *O-*glycosylation in controlling the stability and localization of the model protein FLA4 *in planta*. We found that both intrinsic disorder and glycosylation of the plasma-membrane-linked C-terminal domain can stabilize FLA4 and may act synergistically. This work helps to better understand and rationally design protein stabilizing domains of secreted proteins.

## Introduction

The cell wall of a growing plant cell is a a highly dynamic yet mechanically stable supermolecular composite that undergoes continuous modification through *de novo* deposition and *in muro* remodeling during a plant’s life cycle. While cellulose, hemicellulose and pectin fulfil a variety of biophysical roles, multiple cell wall glycoproteins perform diverse functions, ranging from intercellular signaling and ion exchange to enzymatic and structural roles. However, the functions of their glycans are only partially understood. Protein glycosylations linked to aspartate (*N*-glycans) are structurally and functionally roughly comparable to other eukaryotic systems (Strasser 2016). Like in animals and fungi, plant *N-*glycans act in secretory protein folding and ER-associated protein turnover (ERAD) while the roles of complex *N*-glycans generated in the Golgi apparatus are more diverse but relatively poorly understood (Strasser 2022). Glycosylations linked to hydroxyproline (Hyp) and Ser (*O*-glycans) are unique to the plant kingdom and their structures and functions have evolved independently from *O-*linked glycans in animals and fungi. In fact, plant genomes contain a dazzling multitude of Hyp-rich glycoproteins (HRGPs) including arabinogalactan proteins (AGPs), extensins and Pro-rich proteins (Showalter et al. 2010). The AGP class contains sub-families such as classical AGPs, AG-peptides and chimeric AGPs, all of which contain type II arabinogalactans linked to isolated Hyp residues. Recent biochemical and molecular genetic studies have begun to establish the genes and enzymes involved in type II AG biosynthesis (Strasser et al. 2021), however, the molecular functions of type II AG modifications are only gradually beginning to be understood. Some of the functions have been hypothesized to range from roles in interpolysaccharide crosslinking (Tan et al. 2013; Tan et al. 2023), reversible calcium storage (Lamport et al. 2014; Lopez-Hernandez et al. 2020), to oligosaccharin-like signals (Ma and Johnson 2023) as well as regulating protein fate (Seifert 2020).

A feature of AGPs that contrasts with other HRGPs is their frequent modification by glycosylphosphatidylinositol (GPI)-anchors (Borner et al. 2002; Showalter et al. 2010). In eukaryotes, the GPI-anchor functions as a strong secretion signal and as a determinant for association to plasma membrane lipid raft (Zurzolo and Simons 2016). Moreover, misfolded GPI-APs undergo specialized quality control (QC) mechanisms in the endoplasmic reticulum (ER), at the plasma membrane and the Golgi apparatus (Satpute-Krishnan et al. 2014; Sikorska et al. 2016).

We have previously performed a structure to function analysis of a GPI-anchored chimeric AGP by using different fluorescent protein tagged versions of FASCICLIN-LIKE ARABINOGALACTAN PROTEIN 4 (FLA4) and observed complementation of *sos5/fla4* mutants (Xue et al. 2017). We found that FLA4 was predominantly localized to the plasma membrane and that it was decorated with type II AG recognized by the carbohydrate-specific monoclonal antibodies JIM13 and LM14. Moreover, the combined substitution of six Pro residues located in the hinge region between the two Fas1 domains in combination with the four most highly conserved Pro residues in the C-terminal Pro rich domain resulted in a drastic reduction of cross-reactivity of the mutated FLA4 with AGP-specific antibodies. Moreover, there w a loss of the plasma membrane localization in favor of intracellular compartments co-localizing with endosomal and pre-vacuolar markers. By contrast, a FLA4 variant that lacked the GPI-anchor signal peptide was largely retained in the ER and only to a small proportion reached the apoplast. We proposed that in addition to the GPI-anchor, the type II AG on FLA4 might act either as signal for plasma membrane localization or prevent recruitment into a vacuolar degradation pathway (Seifert 2020; Xue et al. 2017). However, specific structural determinants of FLA4 that support correct localization are presently unknown and the fate of the hypo-glycosylated FLA4 was not clear. To better understand the role of the Pro-rich domains of FLA4 as a model for many GPI-APs that contain AGP-like Pro-rich domains (Borner et al. 2003) and to further elucidate the role of *O*-glycosylation for an individual AGP, we studied additional structural variants in various genetic backgrounds and monitored their cellular fate. We show that substitutions of clustered Pro residues close to the C-terminus with Ala strongly affect normal *in planta* protein stability of FLA4 and cause the redistribution of the protein from the plasma membrane to both the vacuole and the apoplast. Degradation of Pro-to-Ala substitution mutants requires ER to Golgi protein transport, pepstatin A-sensitive proteases and the Golgi-localized secretory regulator ECHIDNA. Surprisingly, *O*-galactosylation of Hyp residues and efficient UDP-D-galactose (UDP-Gal) supply are not essential for normal FLA4 stability. Both ectopic *N*-glycosylation sites as well as *O*-glycosylation motifs partially re-stabilize FLA4 that lacks its normal Pro pattern. The degree of predicted intrinsic disorder at the C-terminus was associated with higher protein stability at the plasma membrane. Remarkably, the substitution of clustered Pro residues with glycosylation preventing but disorder maintaining Gly residues maintained normal FLA4 fate. Taken together, we suggest that while AG II type *O*-glycosylation can contribute to the *in planta* stability of secreted proteins it is intrinsic disorder at the C-terminus of FLA4 that is crucial for *in vivo* protein fate.

## Results

### Substitution of clustered proline residues in the C-terminal domain causes FLA4 destabilization and vacuolar degradation

We previously found that a citrin-FLA4 (F4C) reporter was targeted to the plasma membrane while a mutated version that lacked the six Pro residues in the central Pro rich region and four Pro residues of the eight Pro residues in the C-terminal Pro-rich region was re-distributed to intracellular organelles partially co-localizing with late endosomal markers RabA1e-CFP and RabF2a-CFP (Xue et al. 2017). Both constructs complemented the *sos5* mutant. To more specifically define structural determinants required for FLA4 localization we generated additional Pro-to-Ala substitution variants (supplemental table S1 shows all constructs used in this study). A construct in which only the six central Pro residues in the PR1 domain were replaced by Ala (F4C_P12A) showed plasma membrane localization comparable to ‘wildtype’ FLA4-citrin and some ectopic apoplastic localization (supplemental figure S1). By contrast, a Pro-to-Ala substitution construct targeting the two conserved Pro doublets at the C-terminus (F4C_P34A), showed a loss of the plasma membrane localization and an apoplastic signal comparable to the previously published F4C_P1234A construct (supplemental figure S1). This suggested that structural features in the C-terminal Pro rich region are more critical for FLA4 localization at the plasma membrane than the central Pro residues. We next generated a version lacking all eight Pro residues in the C-terminal region linking the second Fas1 domain and the GPI-modification signal (F4C_P8A). As described previously, F4C localized predominantly at the plasma membrane and also in endosome-like intracellular compartments. The F4C_P8A version, in comparison, showed a pronounced loss of fluorescence signal and a faint endosome-like cytoplasmic signal (supplemental figure S1; Figure 1A). The mCitrin tag on FLA4 is exposed to acidic cellular environments such as the apoplast, the late endosome and the vacuole. It was suggested that the relative pH-sensitivity of mCitrine might be problematic in some situations (Stoddard and Rolland 2019). Hence we generated alternative constructs replacing the mCitrin tag (pKa=5.7) with mTurquoise2 (pKa=3.1) (Goedhart et al. 2012). Like the F4C reporter, the pUBQ10::mTurquoise2-FLA4 (F4T) reporter showed predominant localization at the plasma membrane (Figure 1B, left panel). However, F4T did not label endosome-like organelles but instead, showed a weak diffuse signal in the vacuolar lumen. Like F4C_P8A, the plasma membrane signal of F4T_P8A was strongly reduced compared to the ‘wildtype’ reporter. However, in contrast to F4C_P8A, the F4T_P8A probe showed intracellular localization in the lumen of the vacuole (Figure 1B, left panel). When counterstained with the acidotropic vacuolar marker dye BCEFC-AM and the plasma membrane and endosomal marker dye FM4-64, F4T mostly co-localized with FM4 at the plasma membrane and F4T_P8A co-localized with BCEFC-AM in the lumen of the vacuole (supplemental Figure S2). Accordingly, the correlation of fluorescence signals in root epidermal cells between the FLA4-mTurq2 reporters with the plasma membrane and endosomal reporter dye FM4-64 after short incubation, was around 0.75 with the F4T ‘wildtype’ construct and significantly (P<0.01) lower for the F4T_P8A reporter (Pearsons correlation coefficient R = 0.2) (Figure 1B, right panel).

**Figure 1:**
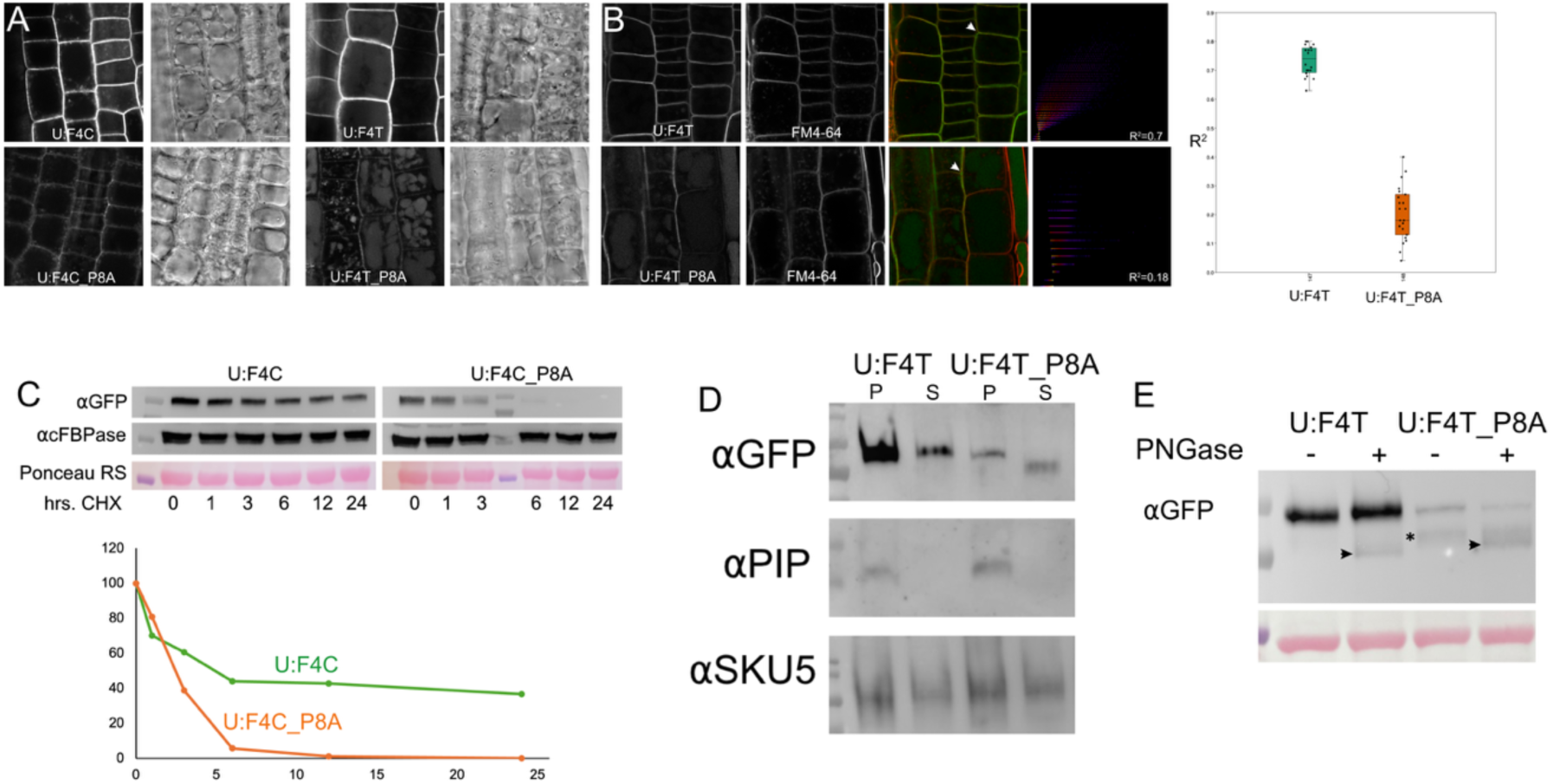
FLA4 is destabilized by Pro to Ala substitutions in C-terminal Pro-rich domain. A) The relatively more pH stable mTurq2 tag shows a vacuolar localization of the F4_P8A variant. B) Wild-type F4T colocalizes with FM4-64 to a higher degree than the F4T_P8A variant. C) Loss of tagged FLA4 versions during CHX mediated block of protein biosynthesis. D) Membrane bound vs. soluble fractions of tagged FLA4 variants. α-PIP antibody is used as control for the microsomal fraction. α-SKU5 stains an abundant GPI-anchored protein. E) PNGase digest of root-extracted tagged FLA4 variants. PNGase sensitive fraction is indicated by arrowhead, biologically unstable fraction is marked by asterisk. Note that in the F4T_P8A variant, PNGase treatment specifically reduces the abundance of the full-length version and not the biologically unstable fraction.

On protein blots of total seedling protein, the signal intensity was generally lower in different F4T_P8A and F4C_P8A transgenic lines compared to the ‘wildtype’ reporter constructs (supplemental Figure S3). As the constructs were all driven by the UBQ10 promoter, possible explanations for the trend of decreased steady state abundance were that either the mutated protein became post-translationally destabilized or translationally suppressed. To distinguish between these explanations, we treated seedlings with the protein biosynthesis inhibitor cycloheximide (CHX) for different times. We monitored reporter abundance relative to total protein loading and the cytosolic housekeeping protein fructose bisphosphatase (cFBPase). The FLA4 reporter proteins underwent a gradual loss compared to total protein and cFBPase, however, while F4C and F4T abundance after 24 hours was around 40% compared to the initial value, the respective F4C_P8A and F4T_P8A variants were strongly reduced already after 3 hours of CHX treatment and became undetectable after 6 hours (Figure 1C). This indicated that the *in planta* lifetime of FLA4 was affected by Pro-to-Ala substitutions in the PR2 domain at its C-terminus.

In addition to its vacuolar signal, the F4T_P8A reporter moreover localized to the apoplast either directly adjacent to the plasma membrane or more brightly to the cell corners where it co-localized with the cell wall reporter EXPANSIN14-mCherry (Samalova et al. 2023) (supplemental Figure S4). This suggests that a fraction of F4T_P8A might be turned over by vacuolar degradation while another fraction is transported to the plasma membrane and released into the apoplast from where it might subsequently diffuse into the culture medium. We next performed fast centrifugation to assess the proportion of membrane-bound to freely soluble reporter protein in relation to the Pro-to-Ala substitutions at the C-terminus. As previously described for F4C (Xue et al. 2017), ‘wildtype’ F4T from roots predominantly partitioned to the membrane fraction (Figure 1D) while F4T_P8A was evenly distributed between membrane-bound and soluble fractions. No apparent difference in molecular weight was observed in the soluble and membrane bound fractions of the ‘wildtype’ reporter. By contrast, the soluble F4T_P8A protein had a smaller apparent molecular weight compared to F4T_P8A in the membrane fraction. Because the tag was attached to the N-terminus of the construct, the reduction in apparent molecular weight suggested that the C-terminus had been proteolytically removed in the P8A variant. To discriminate between FLA4 species before and after the protein had undergone complex *N*-glycosylation in the trans-Golgi we used PNGase F digestion (Strasser 2022). This analysis revealed that bands around 66kDa corresponding to PNGase-sensitive glycoproteins were comparable between the F4C and F4C_P8A reporter lines (Figure 1E, arrowheads), both in apparent molecular weight and intensity. By contrast, the PNGase-resistant fraction of F4T_P8A (Figure 1E, asterisk) revealed lower abundance and decreased apparent molecular weight compared to F4T. This fraction might reflect proteolytic loss of the C-terminus after complex *N*-glycan modification has taken place in the Golgi. Taken together, the replacement of all eight C-terminal Pro residues in the PR2 domain with Ala is accompanied by an almost complete disappearance of FLA4-reporter from the plasma membrane and the concomitant, presumably proteolytic, loss of the C-terminus and release of FLA4-reporter into the apoplast. At the intracellular level, Pro-to-Ala substitution leads to an increased vacuolar localization of FLA4-reporter and an overall acceleration of protein turnover compared to the ‘wildtype’ version. These observations suggested that the ‘wildtype’ and P8A mutant reporters are biosynthesized at comparable rates in the ER but are degraded at markedly different rates after complex glycan attachment in the Golgi with the ‘wildtype’ reporter more stable than the F4T_P8A variant.

*C-terminally mutated FLA4 is degraded in a post-Golgi compartment by aspartate proteases* Using plasma membrane localization in the root and protein abundance after protein biosynthesis inhibition as indicators for *in vivo* protein stability we investigated the possible cellular mechanism involved in the destabilization of the Pro-to-Ala substituted FLA4. To investigate the role of ER to Golgi transport we transformed *F4C* and *F4C_P8A* constructs into the quadruple mutant *p24δ3δ4δ5δ6* (short *p24dquad*) lacking the subclass δ-1 of the p24 cargo receptors and showing a strongly reduced ER to Golgi transport of GPI-APs (Bernat-Silvestre et al. 2020). Similar to the *F4C (sos5-2)* controls, *F4C* (*p24δquad)* was mostly plasma membrane localized, however was additionally in the ER, noticeable by its perinuclear localization. By contrast, *F4C_P8A*(*p24δquad)* was clearly brighter than the *F4C_P8A (sos5)* control and predominantly seen in ER-like structures (Figure 2A). Relative protein abundance after protein biosynthesis inhibition of P8A was apparently higher in the P8A (*p24δquad)* compared to the *F4C_P8A (sos5-2)* control (Figure 2B). PNGase susceptibility of both F4C and *F4C_*P8A was increased in the *p24δquad* compared to the *sos5-2* background confirming increased ER retention of both constructs (Figure 2C). This suggests that normal ER to Golgi transport is rate-limiting for rapid degradation of *F4C_*P8A. This suggestion is consistent with the hypothesis that the Pro-to-Ala mutated FLA4 variants are predominantly degraded in a post-ER compartment.

**Figure 2:**
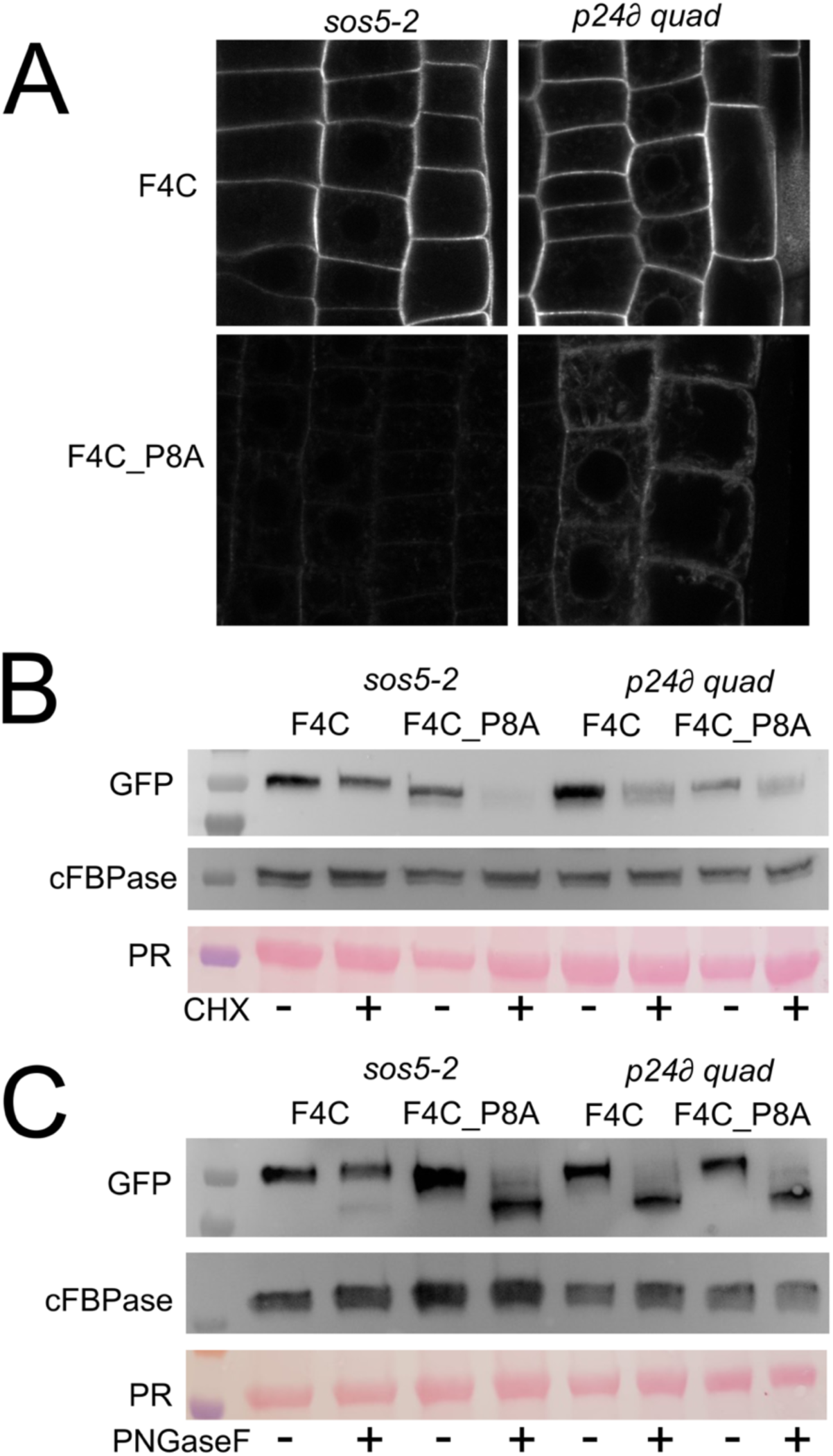
Effect of the transport chaperone p24∂ on the fate of tagged FLA4 variants. A) The ‘wild type’ F4C is marginally ER-like reticulae in *p24∂ quad* mutants. The F4C_P8A variant is stabilized in the ER in *p24∂ quad* compared to *sos5-2*. B) CHX block shows that F4C_P8A is less rapidly degraded in p24∂ quadruple mutants compared to the *sos5* background. C) PNGase digest shows that the *p24∂ quad* mutant background leads to relatively higher ER retention in the of both ‘wild type’ F4C and F4C_P8A. The Ponceau Red (PR) stained Rubisco band and cFBPase are used as loading controls.

Genetic interference with numerous known regulators of post-Golgi protein trafficking failed to reproducibly modulate the destabilization of F4T_P8A (supplemental table S2). However, the loss of function of *ECHIDNA* had an effect on UF4T_P8A trafficking. The ECHIDNA (ECH) protein is localized in the TGN and interacts with YIP4a/b. It acts in the traffic of proteins and cell wall polysaccharides from the TGN to both the plasma membrane and the vacuole (Gendre et al. 2013). The *ech* mutant caused higher brightness of the F4T_P8A probe compared to the isogenic control (Figure 3A) and suppressed the rapid degradation of F4C_P8A in the CHX block experiments (Figure 3B). In epidermal cells in the transition zone between cell division and rapid elongation the *F4T_P8A* (*ech*) probe was seen in bright vacuole-like compartments that were more pleiotropic in shape and more varied in brightness than the vacuoles labelled by *F4T_P8A* (*ECH^+^*) controls. The apoplastic signal also appeared brighter in the *ech* mutants compared to *ECH^+^* wildtype siblings (Figure 3A). The *F4T* (*ech)* lines compared to the *F4T* (*ECH^+^*) isogenic lines showed no difference in *in vivo* protein degradation and microscopically showed comparable signal intensities at the plasma membrane (Figure 3A & B). However, like for F4T_P8A (*ech*), the F4T (*ech*) probe also labelled pleiotropic intracellular structures that were not seen in the *F4T* (*ECH^+^*) controls.

**Figure 3:**
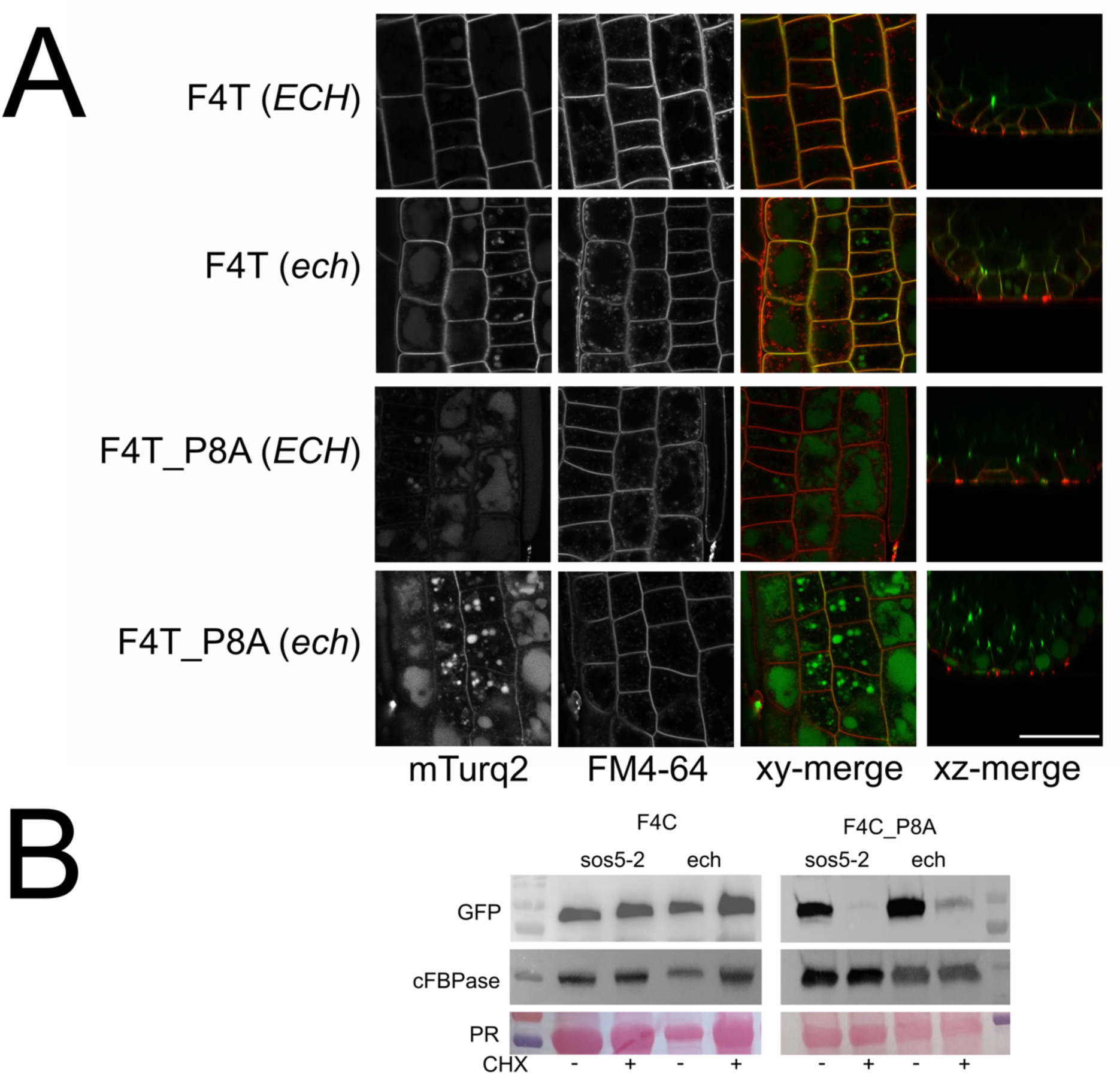
ECHIDNA influences F4T trafficking and the localization and degradation of F4T_P8A. A) Effect of *ech* mutant background on F4T and F4T_P8A localization in roots. B) I*n planta* stability of F4T variants in the *ech* background.

In order to identify the biochemical mechanism leading to degradation of the Pro-to-Ala substituted FLA4 reporters, we took a pharmacological approach and screened different protease inhibitors. DMSO-treated *F4C_P8A* roots showed very little signal at the outer epidermal plasma membrane (supplemental Figure S5 arrowheads). This distribution was not notably affected by leupeptin, AEBSF, MG132, E-64, CP-471474, batimastat or marimastat (all inhibitors at 10µM for 16 h). By contrast, pepstatin A treatment partially restored localization of the F4C_P8A probe at the plasma membrane. The stabilizing effect of pepstatin A was confirmed with the F4T_P8A construct that was seen to clearly co-localize with the plasma membrane after treatment (Figure 4A). Pepstatin A-treated F4C_P8A was also markedly more abundant after cycloheximide treatment compared to DMSO-treated controls (Figure 4B). Taken together, the substitution of potentially *O*-glycosylated Pro residues in the C-terminus of FLA4 leads to destabilization of the protein in a post-Golgi compartment in a process that is sensitive to pepstatin A. Furthermore, the trafficking of both the ‘wildtype’ reporter and the Pro-to-Ala variant is sensitive to mutations in *ECHIDNA*.

**Figure 4:**
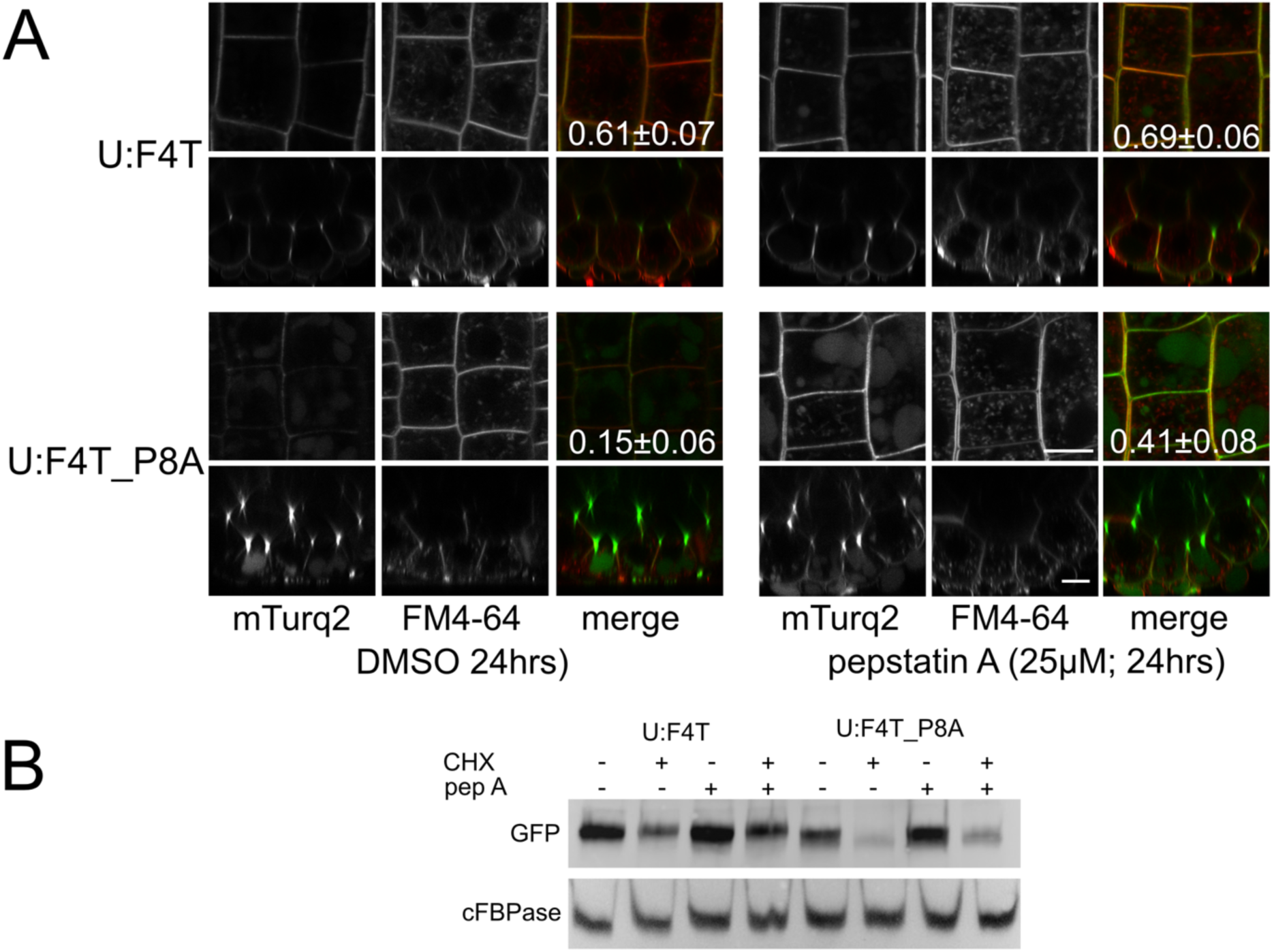
Degradation of F4T_P8A depends on aspartate proteases. A) Pepstatin A stabilizes F4T_P8A at the plasma membrane. B) Pepstatin A increases in planta stability of F4T_P8A

### Proline clusters affect FLA4 stability in synergistic fashion

Substitution of Pro-to-Ala in the C-terminal region of FLA4 results in drastic reduction of protein stability at the plasma membrane in a process that involves post-Golgi trafficking and aspartate proteases. Some Pro residues in this region are potentially *O-*glycosylated but it is not clear whether *O-*glycosylation is a crucial determinant of protein stability. We generated additional mutated variants to better specify the residues that are involved in protein stability. While the combined replacement of the two Pro pairs in position Pro368/Pro367 and Pro389/Pro390 (F4T_P34A) rendered a fusion protein that was as unstable as the F4T_P8A variant, constructs that contained Pro-to-Ala substitutions either in the Pro368/Pro367 position (F4T_P3A) or in the Pro389/Pro390 position (F4T_P4A) showed no notable difference to the ‘wildtype’ construct in localization (Figure 5A) or *in planta* stability (Figure 5B). This observation was not consistent with a specific effect of any one of the two conserved Pro clusters but rather suggested a synergistic effect of both Pro pairs on protein stability.

**Figure 5:**
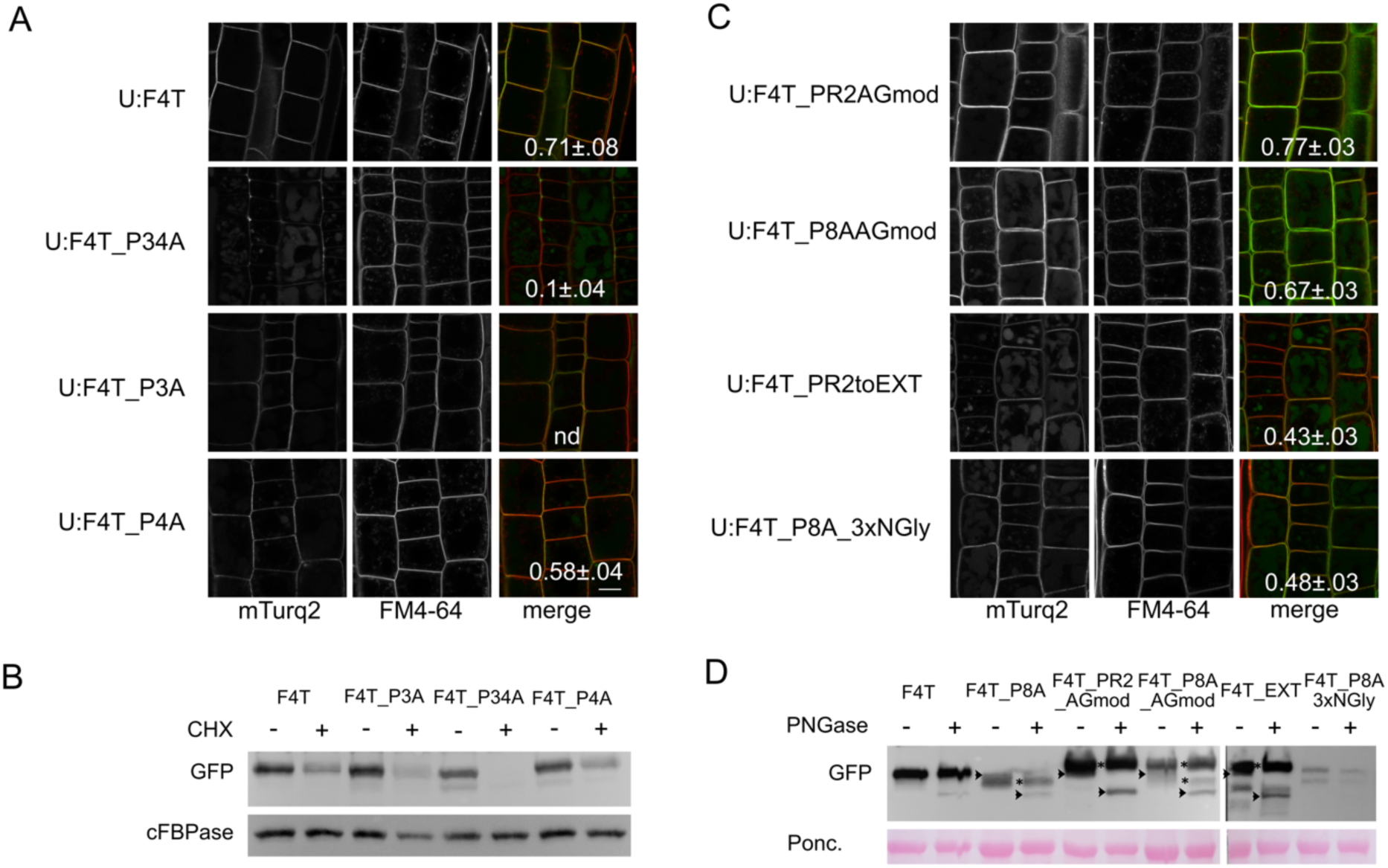
Glycosylation motifs influencing FLA4 fate. A) Pro to Ala substitutions affecting F4T localization. B) Pro to Ala substitutions affecting F4T *in planta* stability. C) Ectopic glycosylation motifs affecting F4T localization. D) PNGase digest of FLA4 constructs containing various glycosylation signals. The arrowheads indicate products that shift by approximately 22kDa and represent *N-*glycosylated protein that carries oligomannosidic *N-*glycan prior to complex *N-*glycan modification and O-glycosylation in the Golgi. Asterisk symbols are placed between respective-PNGase and +PNGase lanes to denote PNGase resistant proteins. These have undergone complex N-glycosylation and in parallel O-glycosylation.

### Diverse protein glycosylation motifs can stabilize FLA4

We next tested whether the loss of Pro residues in their normal position at the FLA4 C-terminus could be compensated for by ectopic motifs known to give rise to *O-*linked or *N-*linked glycans. The *O*-glycosylation module SPSPSP, placed into the otherwise ‘wildtype’ FLA4 C-terminus (*F4T_PR2AGmod*) showed predominant plasma membrane localization (Figure 5C). A tendency of higher abundance of *F4T_PR2AGmod* compared to *F4T* was seen both microscopically and on immunoblots (Figure 5, supplemental figure S6). Compared to the ‘wildtype’ construct, the *F4T_PR2AGmod* variant displayed an increase in apparent molecular weight and appeared as a more ‘smeared’ product, consistent with the SPSPSP sequence being a *bona fide* target for AGP-type *O*-glycosylation. The increased heterogeneity of F4T_PR2AGmod compared to F4T was partly due to the presence of a product band that was sensitive to PNGase F and probably represented an ER-resident protein *en route* to processing in the Golgi (arrowheads in Figure 5D). In F4T, this product was either too low in abundance or too similar in molecular weight with the PNGase-resistant fraction to be discriminated. As mentioned above, the PNGase F resistant fraction is thought to represent the protein that has undergone both complex *N*-glycan processing and *O*-glycosylation in the Golgi.

When the SPSPSP module was added to the otherwise Pro-less FLA4 C-terminus (*F4T_P8A_AGmod*), the reporter protein was predominantly localized at the plasma membrane (Figure 5C) and was as stable as the ‘wildtype’ construct after CHX block (supplemental figure S6). Thus, in the context of a generally Pro-less C-terminus, the presence of the SPSPSP module largely rescues normal protein localization and stability. The PNGase-resistant fraction (asterisks in Figure 5D) consisted of two species with different apparent molecular weights: a major smeary band that migrated as higher molecular weight population compared to F4T and identical to *F4T_PR2AGmod* (upper asterisk in Figure 5D), and a less abundant species that was comparable to the PNGase-resistant fraction of the *F4T_P8A* variant (lower asterisk in Figure 5D). Compared to the *F4T_PR2AGmod,* in the F4T_P8A variant the PNGase-resistant fraction (asterisk in Figure 5D, see also Figure 1E) had a lower molecular weight compared to the PNGase-sensitive (i.e. ER resident) fraction (arrowheads in Figure 5D, see also Figure 1). We did not observe a molecular species consistent with an intact, PNGase-resistant, non-*O*-glycosylated *F4T_P8A_AGmod*. This observation is consistent with the interpretation that the AG module protected the protein from C-terminal degradation only when it was glycosylated.

Replacing the entire C-terminus of FLA4 with a part of the EXTENSIN4 sequence (*F4T_EXT*), yielded a product that localized both in the plasma membrane and the vacuole to a similar extent. This suggested only partial stabilization that contrasted with to the predominantly plasma membrane localized ‘wildtype’ reporter and both constructs carrying the SPSPSP glycomodule. The added EXT module leads to only a minor difference in calculated molecular weight (65.84 for *F4T_EXT* vs 65.39 for *F4T* excluding signal peptides and PTMs), however, it increased the apparent molecular weight of the PNGase-resistant fraction compared to *F4T.* This was not an electrophoretic artefact potentially caused by high Pro-density as the migration of the PNGase-sensitive fraction of F4T_EXT (arrowhead in + lanes in Figure 5D) was similar to all other constructs. The increase in apparent molecular weight of *F4T_EXT* was therefore due to profuse *O*-glycosylation. Compared to *F4T_PR2AGmod*, *F4T_EXT* presented as a more heterogeneous and potentially fragmented product. Therefore, different *bona fide O*-glycosylation modules can overall stabilize FLA4 at the plasma membrane with the AG module being more efficient than the EXT module to prevent vacuolar degradation. This was also observed in CHX block experiments (supplemental figure S6).

Unexpectedly, placing *N-*glycosylation sequons (NXS/T) into the C-terminus of *F4T_P8A*, to generate *F4T_P8A_3xNGly*, also partially stabilized the protein at the plasma membrane (Figure 5C). The electrophoretic analysis showed a major band at the molecular weight of the ‘wildtype’ reporter and a minor band at the molecular weight of the C-terminally truncated fragment that is also formed in the *F4T_P8A* variant (asterisk in the *F4T_P8A lane in* Figure 5D). Despite this partial proteolytic degradation, compared to *F4T_P8A*, *F4T_P8A_3xNGly* had a considerably improved *in planta* lifetime (supplemental figure S6). Taken together, the stability of FLA4 at the plasma membrane can be improved by diverse glycosylation sites with arabinogalactan motifs being more efficient than *N*-glycans and extensin-like sequences.

### Genetic interference with arabinogalactan biosynthesis has no effect on FLA4 stability and localization

Our results indicated a synergistic role of clustered, potentially *O*-glycosylated Pro residues for protein fate that can be compensated by a variety of glycosylation motifs. To further test whether type type II AG *O-*glycosylation is critical for FLA4 protein fate we used genetic tools to alter glycan biosynthesis and tested effects on FLA4 protein fate. The suppression of UDP-Gal biosynthesis in mutants defective in UDP-glucose 4-epimerase (UGE) was previously shown to affect the galactose content in the cell wall and to affect composition of xyloglucan, pectin and AGPs (Seifert et al. 2002; Rösti et al. 2007; Andeme-Onzighi et al. 2002; Nguema-Ona et al. 2006). Surprisingly, the *uge2 uge4* double mutant, which was previously shown to display abnormal growth and cell wall galactose content in roots and whole seedlings (Rösti et al. 2007) did not notably affect F4T localization in the root epidermis (Figure 6A). Likewise, the *in vivo* steady state abundance and *in planta* stability of F4T (*uge2 uge4*) was unaffected compared to the isoparental *F4T* (*sos5)* control (Figure 6A). The *galt2 galt5* double mutant and the *hpgt1 hpgt2 hpgt3* triple mutant were previously shown to drastically affect AGP-biosynthesis (Ogawa-Ohnishi and Matsubayashi 2015; Basu et al. 2015) and to act in a linear genetic pathway with *FLA4/SOS5* with respect to all observable phenotypes (Showalter and Basu 2016). However, neither mutant background notably affected F4T localization (Figure 6A), abundance and stability (Figure 6B).

**Figure 6:**
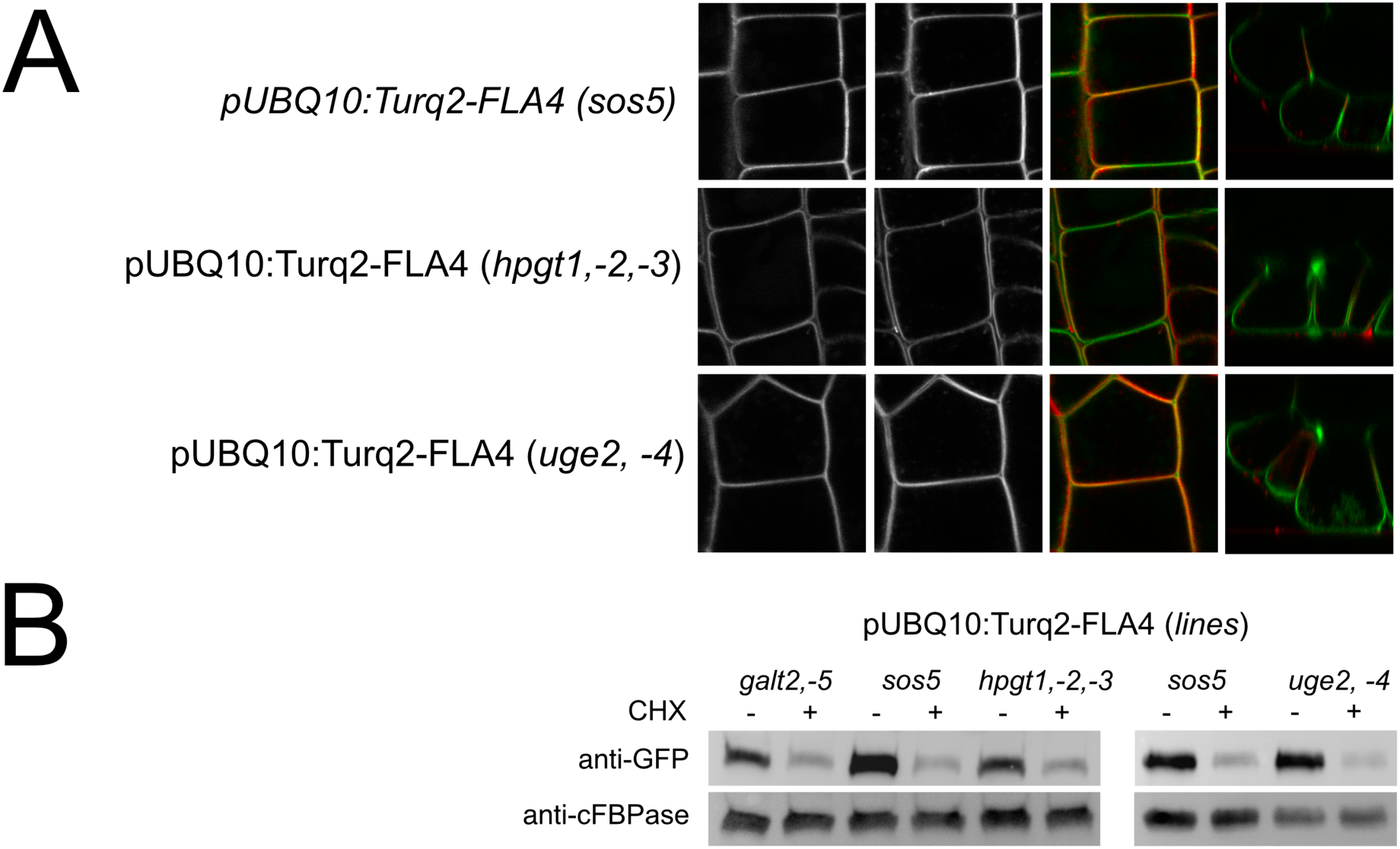
Genetic interference with O-glycosylation does not affect FLA4 fate. A) Localization of F4T in the root epidermis in *sos5-2*, *hpgt1 hpgt2 hpgt3* and *uge2 uge4* mutant backgrounds. B) *In planta* protein stability is comparable in *sos5-2*, galt2 galt5 *hpgt1 hpgt2 hpgt3* and *uge2 uge4* mutant backgrounds.

### Proline to Alanine substitutions affect intrinsic disorder and secondary structure

So far, we found that diverse structural interventions affecting glycosylation of the C-terminus influenced FLA4 protein fate with a clear trend that both *O*-linked and *N*-linked glycosylation increases protein stability. This contrasted with the lack of effect on FLA4 fate by genetic suppression of Hyp *O-*galactosylation. One possible explanation for this discrepancy might be that, in addition to the prevention of *O*-glycosylation, the introduction of Pro-to-Ala substitutions might have led to other structural alterations as a cause for protein destabilization.

To better understand this inconsistency, we modelled the structures of wildtype and various mutated forms of the Pro rich C-terminal region using computational tools. Regions rich in Pro, Ala, Ser and Thr (PAST) are a typical indicator of Pro-4-hydroxylation and *O*-glycosylation in HRGPs (Showalter et al. 2010). On the other hand, it has been noted that PAST regions display high intrinsic disorder (Shafee et al. 2020; Johnson et al. 2017a). Intrinsically disordered regions do not fold into well-defined structures and are enriched in amino acids Ala, Arg, Gly, Gln, Ser, Pro, Glu and Lys, and depleted in Trp, Cys, Phe, Ile, Tyr, Val, Leu, Asn (Xiao and Xia 2025; Williams et al. 2001; Theillet et al. 2013). Based on these compositional characteristics, the Pro-to-Ala substitutions were not expected to affect intrinsic disorder to a large degree. Nonetheless we determined intrinsic disorder using prediction tools. Most of the FLA4 wildtype protein shows intrinsic disorder values well below the 0.5 threshold (Figure 7A), except for the C-terminal Pro-rich domain that had an average AIUPred value of 0.89. Unexpectedly, the Pro-to-Ala substitutions greatly affected intrinsic disorder prediction for various experimentally tested variants. For instance, the P8A, P6A and P34A variant displayed much lower than wildtype intrinsic disorder in the C-terminus with average AIUPred value of 0.23, 0.45 and 0.61 (Figure 7B), respectively, which roughly mirrored their effect on *in planta* protein stability and localization. By contrast, variants F4T_P3A and F4T_P4A that showed a moderate effect on protein stability presented relatively high AIUPred profiles with values of 0.81 and 0.8, respectively (Figure 7B). These results suggested a positive correlation between intrinsic disorder of the FLA4 C-terminus and protein stability. The introduction of glycosylation motifs into the C-terminus was associated with increased protein stability. This was not entirely reflected in AIUPred values. While the variant PR2_AGmod that was relatively more stable compared to the wildtype, also presented a higher AIUPred value of 0.91, the stable F4T_P8A_AGmod variant only presented a AIUPred of 0.4 that was only a minor recovery compared to F4T_P8A (0.23), and was also below both the biologically unstable F4T_P6A and F4T_P34A versions. Also, for the *N*-linked glycan variant P8A_3xNGly, the AIUPred average value of 0.48 was below the value of F4T_P34A. These comparisons should be interpreted with caution because currently available disorder-prediction tools do not account for the effects of protein glycosylation.

**Figure 7:**
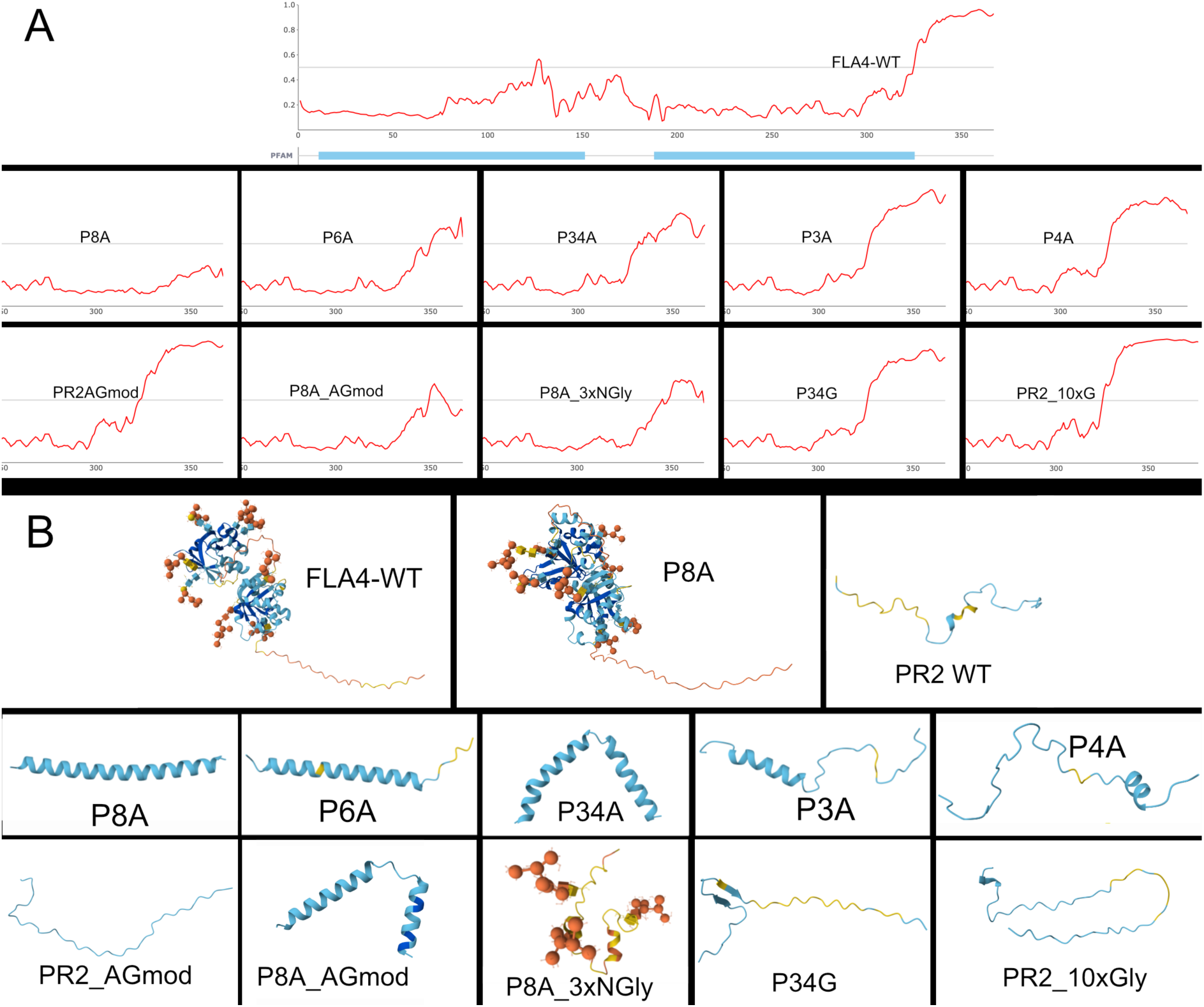
Structure predictions of different FLA4 variants used in this study. A) Intrinsic disorder profile of FLA4 predicted by AIUpred. The profiles of different variants are only shown in the C-terminal domain. Note that AIUpred cannot predict the effect of protein glycosylation on intrinsic disorder. B) Alphafold3 prediction of FLA4 and FLA4_P8A modeled as whole proteins without the N-terminal secretion peptide and the C-terminal GPI-signal. The C-termini of the wild type and the different variants are subsequently modeled in isolation. Note that the C-terminal region in the whole model of the P8A variant is disordered but shows an alpha helix with high confidence when modeled in isolation. Note that the O-glycosylation introduced in the PR2_AGmod and the P8A_AG versions cannot be predicted by AIUpred or Alphafold3 server and the effect of O-glycosylation in protein disorder is presently unknown.

An Alphafold 3 prediction of the FLA4 wildtype protein that included *N-*linked glycans showed a pair of two well defined globular Fas1 domains abutted by an unstructured C-terminal domain and the P8A variant presented a similar structure (Figure 7B). However, modeling the C-terminal domains in isolation revealed a different picture that was in good agreement with the AIUPred profile. The wildtype model essentially lacked secondary structure. By contrast, the P8A variant of the C-terminus was predicted to adopt a predominantly α-helical secondary structure (Figure 7B). Different Pro-to-Ala variants showed a varying degree of α−helix secondary structure that negatively correlated with intrinsic disorder and *in planta* protein stability. The exception to this trend was the F4T_P8A_AGmod construct that contained a SPSPSP module in an otherwise Pro-to-Ala mutated C-terminus and that showed good stability *in planta* but in addition to relatively low intrinsic disorder a largely α-helical secondary structure. By contrast the version F4T_P8A_3xNGly modeled with *N-*linked glycans was largely free of defined secondary structure (Figure 7B). The peptide without *N*-glycans modeled into a largely α-helical structure (supplemental Figure S7).

### Clustered proline to glycine substitutions preserve both intrinsic disorder and protein fate

The unexpected prediction that Pro-to-Ala substitutions might lead to an α-helical secondary structure replacing of an intrinsically disordered domain, together with our experimental observations that were inconsistent with an exclusive role of specific *O*-glycans for protein stability, led us to speculate that not necessarily *O*-glycosylation but intrinsic disorder might be crucial for FLA4 stability. We modeled the structural effect of alternatives for Ala as substituted residue. We found that in the Pro-to-Gly variant P34G, intrinsic disorder was much higher than in the P34A variant (0.822 and 0.611, respectively). More clearly, in strong contrast to P34A that formed α-helices, the prediction for the P34G variant was devoid of secondary structure (Figure 7). A variant that increased predicted intrinsic disorder (0.939) compared to the wildtype contained a 10-residue Gly linker between the two Pro clusters (PR2_10xG). Both versions were transformed into Arabidopsis plants.

Consistent with the hypothesis that intrinsic disorder in the C-terminal domain of FLA4 is crucial for protein stability, we observed that the *F4T_P34G* variant showed a predominantly plasma membrane localization similar to the *F4T* ‘wildtype’ and in dramatic contrast to F4T_P8A and F4T_P34A (Figure 8A). Likewise, when *in planta* protein stability was assayed using CHX, the stability of the F4T_P34G reporter was comparable to the F4T ‘wildtype’ variant (Figure 8B). The artificially increased intrinsic disorder in the PR2_10xG variant, did not significantly alter stability or localization of FLA4 (supplemental figure 8). In summary, FLA4 is stabilized *in planta* by its intrinsically disordered C-terminus. Both amino acid composition and glycosylation contribute to intrinsic disorder and protein fate.

**Figure 8:**
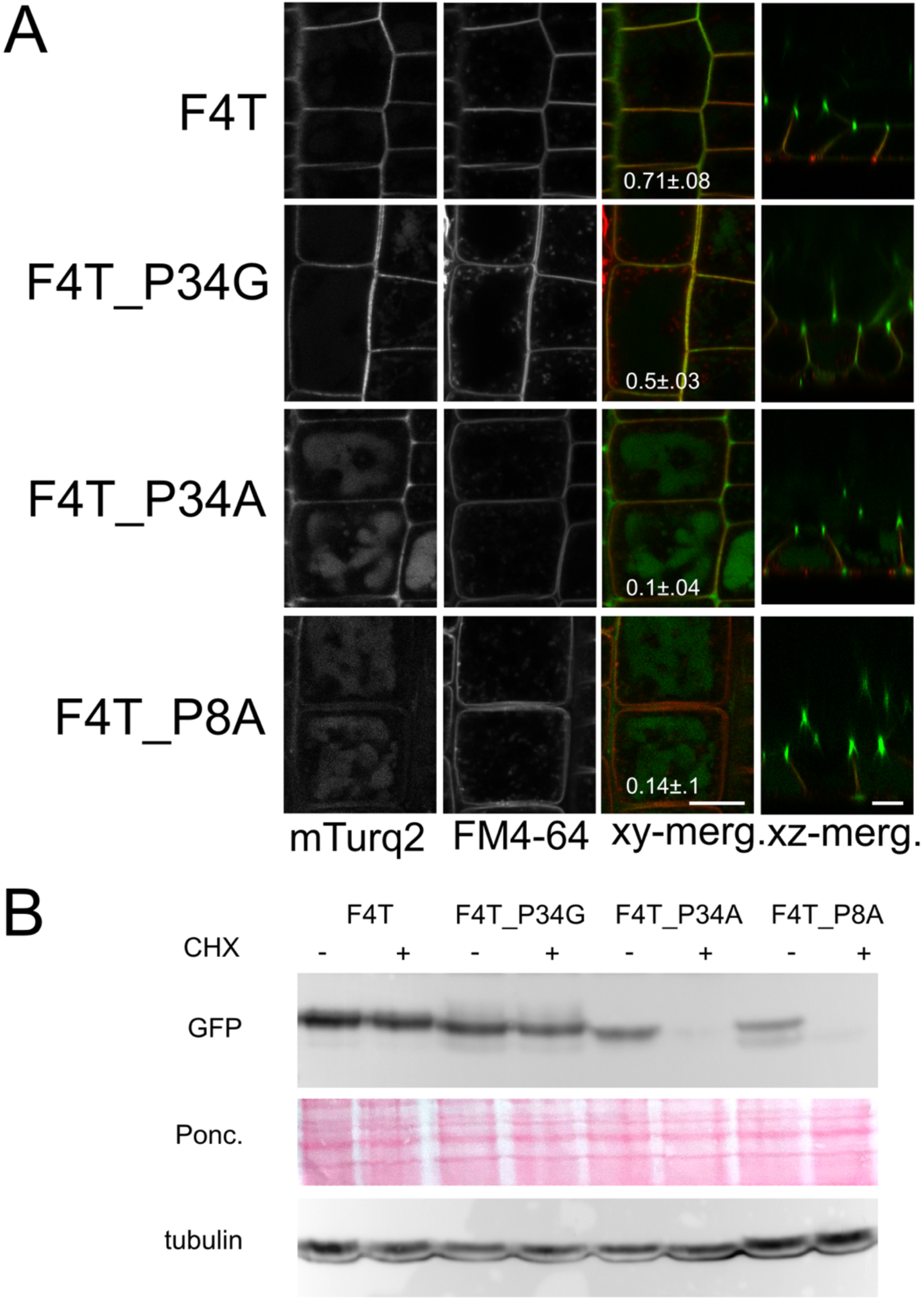
Pro to Gly substitution in *O-*glycosylation sites conserve normal protein fate. A) The F4T_P34G variant is mostly localized to the plasma membrane like the ‘wild type’ F4T, compared to the vacuolar localization of the F4T_P34A and the F4T_P8A variants. B) *In planta* protein stability in the F4T_P34G variant is similar to ‘wild type’ F4T and clearly higher than the F4T_P34A and the F4T_P8A variants.

## Discussion

In this study we investigated the potential role of *O*-glycosylation on the cellular fate of FLA4 as a model for type II AG modified GPI-anchored proteins. Previous observations indicated a role of type II AG glycosylation for the fate of various glycoproteins (Ma et al. 2022; Ma et al. 2023; Teh et al. 2022; Borassi et al. 2020; Seifert 2020) including FLA4 (Xue et al. 2017). In the present study, we found that the interplay of intrinsic protein disorder and *O-*glycosylation motifs that was observed on a genome-wide scale (Shafee et al. 2020; Johnson et al. 2017a) was crucial for normal protein fate of FLA4. The reduction of intrinsic disorder in the C-terminal region of FLA4 lead to its proteolytic degradation by aspartyl proteases in a late secretory compartment. The functional investigation of *O-*glycosylation in plants is encumbered by the relative lack of precision in predicting and detecting hydroxylated and *O-*glycosylated Pro residues in a secretory protein. An important rule called the Hyp-contiguity hypothesis postulated that stretches of contiguous Hyp residues are arabinosylated giving rise to extensin-linked short arabinosides and that clusters of isolated Hyp residues are galactosylated and form the basis of more extensive type II AGs (Shpak et al. 2001; Kieliszewski 2001). However, variations in the Pro-hydroxylation and Hyp-glycosylation of different artificial HRGPs suggested additional significance of flanking amino acids (Tan et al. 2003). Most previously characterized AGPs contain stretches of directly adjacent repetitive (XaaPro) units, and plant-wide studies have used a rule of at least three [(Ser/Thr/Ala/Val)Pro] motifs in a ten residue sequence window as defining criterion for AG-domains (Johnson et al. 2017b; Johnson et al. 2017a; Shafee et al. 2020). This is not the case for FLA4 that contains three [[(Ser/Thr/Ala/Val)Pro] motifs that are each separated by 7 and 12 intervening residues, respectively. Nevertheless, FLA4 has been shown to carry type II AG recognized by the monoclonal antibodies JIM13 and the LM14 that were suppressed by Pro-to-Ala substitution of clustered Pro residues which led to mislocalization to the apoplast and pre-vacuolar compartment of FLA4-reporter (Xue et al. 2017). As we noted in the course of this study the Pro-to-Ala substitutions came at the price of interfering with intrinsic protein disorder.

The structure to function analysis of individual amino acid residues by site directed mutagenesis often uses Ala as substituted residue - a strategy commonly referred to as ‘alanine scanning’ (Kokoszka and Kay 2015; Morrison and Weiss 2001; Sadeghi and Birnbaumer 1999; Cunningham and Wells 1989). The main reasons for choosing Ala are the absence of a side chain beyond the β-carbon, minimal impact on main chain conformation (which is a potential consequence when using Gly as substituted residue) and Ala being the most abundant amino acid that is found in both buried and exposed positions (Cunningham and Wells 1989). Other amino acids are preferred in specific cases such as Gln as the obvious choice to replace Asn in the study of *N*-glycosylation motifs AsnXaa[Ser/Thr]. There are some examples of ‘alanine scanning’ studies to assess the role of *O*-glycosylation in HRGPs that showed that mutation of the Pro-rich region of various AGPs and FLAs interfered with normal function, localization and protein abundance (Ma et al. 2023; Borassi et al. 2020; Xue et al. 2017; Teh et al. 2022). While focusing on one member of the AGP superfamily, the present study provides a note of caution to the application of the ‘alanine scanning’ strategy for the study of HRGPs.

An approach to study AGP glycan function that is complementary to site specific mutagenesis, utilizes genetic interference with glycan biosynthesis by introducing mutations in type II AG glycan-specific glycosyl transferases or, less specifically, mutants defective in nucleotide sugar metabolism and transport. Combined mutations in Hyp-specific galactosyl transferases (GALTs) were found to strongly reduce the abundance of AGPs (Ogawa-Ohnishi and Matsubayashi 2015; Basu et al. 2015; Kaur et al. 2023) providing genetic tools to study the roles of AGP glycosylation. Interestingly, the GALT double mutants *galt2 galt5* and triple mutant *hpgt1 hpgt2 hpgt3* phenocopied the root and seed coat phenotypes of *sos5* indicating a role of these GALTs in *FLA4* function (Showalter and Basu 2016). Moreover, the *uge4* single and the *uge2 uge4* double mutants defective in the supply of UDP-D-galactose were previously associated with alterations of AGPs in roots and whole plants, respectively (Ding and Zhu 1997; Andeme-Onzighi et al. 2002; Seifert et al. 2002; Rösti et al. 2007). Surprisingly, neither localization nor *in planta* lifetime of FLA4 reporters were notably affected by the *hpgt1 hpgt2 hpgt3*, *galt2 galt5*, or *uge2 uge4* mutant combinations. In principle, the absence of any effects of reduced *O*-glycosylation on FLA4 might be due to genetic redundancy as more than 30 genes encoding putative AGP-specific GALTs exist in the Arabidopsis genome (Qu et al. 2008). Indeed, recently generated quintuple and octuple mutants showed slightly enhanced growth defects compared to *galt2 galt5* and *hpgt1 hpgt2 hpgt3* (Kaur et al. 2023). Likewise, the *uge1 uge2 uge4* triple mutant showed a more severe reduction of growth than *uge2 uge4* (Rösti et al. 2007). The alternative explanation is that not only type II AG modification of the PR2 domain but also other structural features that are crucial for protein fate were disturbed by the Pro-to-Ala substitutions. This suspicion was confirmed by analyzing the *F4T_P34G* variant that lacked AG modification sites but showed normal protein localization and stability. As opposed to *F4T_P34A,* this construct was predicted to maintain high intrinsic disorder in the C-terminal domain.

Hence, we observed an overall positive relation between predicted intrinsic disorder in the C-terminus adjacent to the GPI-anchor and stability of FLA4 and a negative relation between predicted α-helical secondary structure and stability. While this finding was unexpected, the high thermal and environmental stability of intrinsically disordered proteins (IDPs) and intrinsically disordered protein regions (IDPRs) has previously been highlighted in a commentary by (Uversky 2017). In this respect it is noteworthy that both Ala and Gly, while showing quite contrasting effects in the P34A and the P34G constructs, respectively, have a very similar statistical distribution between ordered and disordered proteins and are generally regarded as mildly disorder promoting (Xiao and Xia 2025; Williams et al. 2001; Theillet et al. 2013). Nonetheless, both AlphaFold 3 and AIUPred highlighted the differential effects of these substitutions on predicted disorder in the FLA4 C-terminus. One notable exception to the trend that intrinsic disorder correlated with protein stability was the *F4T_P8A_AGmod* construct that showed a relatively low ID and mostly α-helical structure but was even more stable than the ‘wildtype’ construct. The increased mass and mass heterogeneity observed in FLA4 variants modified with an SPSPSP motif in its PR2 domain (F4T*_AGmod* and *F4T_P8A_AGmod*), showed that these were more highly glycosylated than the ‘wildtype’. On the one hand this observation underlines the notion that Pro hydroxylation and glycosylation does not follow simple sequence rules but depends on the structural context. On the other hand, if our hypothesis that intrinsic disorder at the C-terminus is essential for protein stability is correct, it might mean that the introduction of the SPSPSP module might increase protein disorder not only through the introduction of disorder-promoting Ser and Pro residues but also through the addition of large type-II AG modifications. At present this suggestion is speculative as no modelling or molecular dynamics simulation of the folding behavior of AGP-like proteins are known to the authors. Compared to FLA4 ‘wildtype’ constructs and even more strikingly compared to constructs containing the ectopic AG-module, the replacement of the FLA4 C-terminus with a repetitive section of *EXT4* (*F4T_PR2toEXT*) led to fragmentation of the resulting protein and its release into vacuole and apoplast. Our observation was consistent with previous studies with AG-glycomodules or extensin-domains coupled to GFP (Zhang et al. 2019), and it confirms that type II AG modifications are superior to extensin domains with respect to protein stabilization. In the light of our study that suggests a relation between protein intrinsic disorder and biological stability and the notion that the arabinosylated extensin domains are regarded as biophysically rigid (Velasquez et al. 2015; Stafstrom and Staehelin 1986; Heckman et al. 1988), a disordered and conformationally unconstrained protein domain might be an essential ingredient of high protein stability. Compared to the biologically unstable *F4T_P8A* variant, the incorporation of three *N*-glycosylation sites partially stabilized the *F4T_P8A_3xNGly* product at the plasma membrane. The relation between protein *N-*glycosylation and disorder is complex. On the one hand, *N-*glycans have a stabilizing effect on the formation of well-structured protein domains and are an integral component of the ER-related protein degradation system (ERAD) (Hao et al. 2025) both which is consistent with the notion that *N*-glycans might act towards the formation of properly folded protein (domains). On the other hand it was suggested that *N-*glycans once attached to the protein behave in a highly dynamic way thereby generating intrinsically disordered regions at the surface of glycoproteins (Gazaway et al. 2025). Our observations are consistent with *N-*glycans increasing disorder in the α-helical structure that was generated with the P8A mutation, and with that, increasing stability of FLA4 at the plasma membrane. However, we also noted that several independent transformant lines expressing P8A_3xNGly under the control of the UBQ10 promoter, showed relatively low steady state content of F4T_ P8A_3xNGly protein compared to most other constructs. It is possible that the positioning of the *N-*glycosylation sites in the C-terminus might somehow reduce protein biosynthesis. This notion is compatible with the co-translational nature of *N-*glycosylation. Furthermore, it is interesting to note that classical AGPs and AG-peptides that consist of stretches of discontiguous Hyp, are never *N*-glycosylated. By contrast chimeric AGPs that in addition to their Hyp-rich, intrinsically disordered PAST domains, also contain well-ordered globular domains such as Fas1 domains (Johnson et al. 2003), Phycocyanin domains (Poon et al. 2012) or PAC domains (Hijazi et al. 2012) and it is common that those carry Asn-Xaa-Ser/Thr sequons for *N*-linked glycosylation. Given their role in protein folding control, the co-translational addition of *N*-glycans to IDRs might slow down protein biosynthesis or even spuriously trigger ERAD. In this respect it is interesting to note that in eukaryotic genomes including plants there is a high correlation between intrinsic disorder and *O*-glycosylation but no correlation or even an inverse relationship between intrinsic disorder and *N*-glycosylation (Kurotani et al. 2014).

This could mean that in the ER the IDRs of AGPs are only modified by prolyl-4-hydroxylation and potentially by the transfer of initial galactoses. *O*-glycans are mainly built up in the Golgi at which stage they might contribute to protein stability not by promoting folding but by supporting intrinsic disorder. This hypothesis is currently not straightforward to test directly. However, future bioinformatic studies to model protein folding and disorder with respect to Hyp-linked *O-*glycosylation, would not only help to better interpret glycoprotein fate and function but also help to rationally design stability-modulating tags in recombinant protein production.

Irrespective of the precise structural requirement for FLA4 *in planta* fate, F4T_P8A showed a dramatic suppression of *in planta* lifetime and plasma membrane localization. We investigated the genetic and biochemical effectors of the cellular fate of the destabilized FLA4 variant. The route of both the normally glycosylated wildtype FLA4 and the F4T_P8A variant is assumed to follow the classical secretory pathway with the participation of the ER to Golgi cargo receptors of the p24∂ family that were previously shown crucial for transporting GPI-anchored proteins(Bernat-Silvestre et al. 2020). As expected we found that p24∂ isoforms facilitate the efficient ER exit of both F4C variants which was rate-limiting for the rapid degradation of F4C_P8A. In the ER, FLA4 first is modified co-translationally with presumably eight *N-*glycans, receives its GPI anchor by the transamidase GPI8 and some of its C-terminal Pro residues are hydroxylated. The transamidase reaction is thought to occur strictly after full synthesis and *N*-glycosylation of the protein but does not depend on prior protein folding (Spurway et al. 2001). Therefore, we think that in the ER, F4T and its Pro-to-Ala variants behave in a comparable fashion. Consistently we have not observed an effect on F4T_P8A lifetime of the *mns4 mns5* mutant (supplemental table 2) that suppresses ER-dependent protein degradation (Huttner et al. 2014). During processing in the Golgi, *N*-glycans become PNGase-resistant by obtaining complex modifications. Hyp-linked type II AGs are attached to FLA4; these glycans are not large enough to cause a clear shift of apparent molecular weight but are sufficiently complex to be recognized by LM14 and JIM13 (Xue et al. 2017). Upon Golgi exit FLA4 passes the TGN from where it is mostly targeted to the plasma membrane. We assume that the TGN is a site where glycoprotein integrity is checked and misfolded cargo is redirected to degradation in the lytic vacuole. Indeed, some Pro-to-Ala mutated constructs are observed in the vacuole. This vacuolar localization seems to be triggered primarily by the abnormal structure of mutated FLA4 variants but not necessarily by abnormal *O-*glycosylation, because we did not see vacuolar FLA4 in *O-*glycan defective mutant backgrounds. The strongly reduced *in planta* stability of F4T_P8A partially depends on TGN function as indicated by the higher apparent stability in the *ech* mutant background. This dependence could be due to the strong disruption of TGN mediated transport in *ech* which is know to affect trafficking of multiple membrane proteins and polysaccharides (Gendre et al. 2011; Boutte et al. 2013; Liu et al. 2023). While it has been hypothesized that *ECH* is particularly important for the secretion of cell wall polysaccharides we did not observe a differential effect of *ech* on the wildtype or the glycan-defective form of FLA4. However, the increased *in planta* protein stability of F4T_P8A in the *ech* background compared to *ECH* wildtype siblings might be the result of inefficient transport to the plasma membrane that might be the major site of proteolytic degradation of F4T_P8A, potentially executed by apoplastic proteases. A family of proteases potentially involved in degradation of FLA4_P8A are aspartate proteases sensitive to pepstatin A. Among hundreds of proteases in Arabidopsis, there are at least 60 aspartyl proteases (van der Hoorn 2008). This protease family also contains about ten members predicted to be GPI-anchored (Borner et al. 2003). Interestingly, two members of this family named A36 and A39, that are preferentially expressed in germinating pollen are required for normal cell wall composition in pollen (Gao et al. 2017). Other proteases of this family that are more generally expressed seem potential candidates involved in the degradation of FLA4 or its variants that might be involved in cell wall modification. We also found that mutations in *ech* affected the localization of wildtype FLA4-reporter and both degradation and localization of the F4T_P8A variant. Because *ECH* is involved in the trafficking of many plasma membrane and extracellular proteins and polysaccharides, the effect of *ech* on the degradation of F4T_P8A might be due to abnormal trafficking of GPI-anchored aspartyl-proteases to the plasma membrane. By contrast, mutations in many known regulators of vacuolar degradation did not notably affect the degradation of F4T_P8A. Hence, one possible scenario is that only a small fraction of Pro-to-Ala mutated FLA4 is targeted to the vacuole for degradation while most of it is transported to plasma membrane where its abnormal C-terminus is recognized by GPI-anchored aspartyl proteases that proteolytically shed the protein into the apoplast. The rapid loss of the signal in the CHX block experiment might be due to diffusion of the soluble protein from the seedlings into the culture medium rather than processive proteolysis. Consistently, F4_P8A was detected more abundantly in the apoplastic fluid than in the cellular fraction, compared with the ‘wild-type’ FLA4 reporter.

### Conclusions

In summary, we suggest that the role of type II AG *O-*glycans for protein fate is tightly interrelated with intrinsic protein disorder. Given the importance of intrinsic disorder for biomolecular condensates in all cellular compartments including the apoplast (Miao et al. 2026) this relation warrants further biochemical and bioinformatic investigation.

## Methods and Materials

### Plant materials and Constructs

*Arabidopsis thaliana* seeds were grown on plates as previously described (Xue et al. 2017). Stable transformation of various FLA4 constructs was performed as previously described into *sos5-2* mutants (Xu et al. 2008).

All constructs were based on pUBQ10:FLA4-citrin (Xue et al. 2017). To generate pUBQ10:mTurq2-FLA4 (F4T) the sequence of mTurquoise2 (Goedhart et al. 2012) was amplified using appropriate primers to replace mCitrin. Fragments containing different amino acid substitutions were chemically synthesized and cloned into the EcoRI/NotI fragment of F4T by a commercial service (BioCat, Heidelberg). Wildtype and mutant peptide sequences and their abbreviations are provided in supplemental Table S1.

### Chemical treatments

For cycloheximide (CHX) treatments, seeds were germinated in liquid half strength Murashige & Skoog medium (salts and vitamins; Duchefa) and 1% sucrose (MS/2). 5-day-old seedlings were harvested by wicking off liquid on paper tissue, weighing and briefly storing on ice before extraction in 2-mL plastic tubes. For protein extraction, two steel balls were added and samples were ground in a swing-mill (MM 2000; Retsch, Haan Germany) at 30 min^-1^. for 2 min. Extraction buffer (Tris/HCl 50 mM pH8, EDTA 20 mM, DTT 1 mM, 0.5% Triton X-100, 1% protease inhibitor cocktail [Merck, Austria]) was added at a 3:1 ratio (µL/mg fresh weight) and incubated on ice for 10 min. To clear cellular debris, the homogenate was centrifuged twice at 13000 x g for 10 min. Treatment with CHX (final conc. 180µM) was for 3 h or for the indicated times at room temperature in ambient light and temperature conditions. Protease inhibitors used were leupeptin, AEBSF, pepstatin A (all from Lactan, Graz), MG132, E-64, batimastat, marimastat, CP-471474 (all from Merck, Austria) and prepared as 20mM stocks in DMSO. Digestion with PNGase F (New England Biolabs) was performed according to manufacturer’s instructions as previously described (Huttner et al. 2012). Microsomal and soluble proteins were fractionated as previously described (Xue et al. 2017), except that centrifugation was in a 100µl volume at 70.000 x g for 60min.

### Microscopy

FM4-64, a styryl dye labelling the plasma membrane and upon prolonged exposure the membranes of the endosomal pathway and the tonoplast, was added from a DMSO stock solution to a final concentration of 0.5 mg L^-1^ in MS/2 medium and seedlings were incubated for 10 minutes before observation in dye-free medium. Under these conditions the FM4-64 signal was found predominantly at the plasma membrane and to a lesser extent at the early endosomes. Images were acquired with a Leica Plan Apo 63/1.2 water immersion lens at x4 zoom (pixel dimension 60 nm^2^) in overlapping frames focused at the root epidermis from the meristem to the early elongation zone. To acquire two-channel images using confocal laser scanning microscopy (CLSM), the mTurq probe and the FM4-64 dye were simultaneously excited at 458nm and 514nm, respectively and signals were sampled at 470 to 505 nm and 590 to 680 nm respectively. To obtain a measure of co-localization between the two probes, the Coloc 2 plugin in the FIJI freeware was used and Pearson’s R (no threshold) was recorded. To label vacuoles 2’,7’-Bis-(2-Carboxyethyl)-5-(and-6)-Carboxyfluorescein, Acetoxymethyl Ester (BCEFC-AM, Thermofisher Austria) was used at a final concentration of 1.6µM to stain seedlings for 20 minutes and recorded at Ex=488nm and Em=510-540 in sequential mode after mTurq2/FM4-64.

### Western Blotting

Protein electrophoresis, blotting and detection of GFP was performed as previously described (Xue et al. 2017). We used Ponceau Red (Lactan, Austria), rabbit anti-cytosolic Fructose Bisphosphatase (cFBPase; Agrisera, Sweden), or rat-anti α-tubulin (YOL1/34, Thermofisher, Austria) as loading control and reference in determining relative signals on Western blots as shown previously (He et al. 2023). For relative quantitative comparison, we used the *Analyze*/*Gels* tool from FIJI to determine band intensity. Microsomal fractions were re-probed with rabbit anti-PIP (Agrisera, Sweden) and rabbit anti-SKU5 antiserum (Sedbrook et al. 2002).

### Bioinformatics

Intrinsic disorder prediction was performed using the *AIUPred* sever (Erdos et al. 2025; Erdos and Dosztanyi 2024)(https://iupred.elte.hu) that combines high predictive power with fast computation time (Erdos and Dosztanyi 2024). The FLA4 peptide sequence from I29 to S396 and its engineered variants were used for the prediction of the *AIUPred-Only disorder* value with *default smoothing*. To obtain the numerical values for intrinsic disorder in the C-terminal region that were described in the text, the average prediction values between P358 to S396 were calculated. Alphafold 3 predictions were performed with default parameters (Abramson et al. 2024). When *N*-linked glycans were modeled, we generally used a hexamannosidic glycan with the notation NAG(NAG(MAN(MAN(MAN)(MAN(MAN)(MAN))))).

### Artificial Intelligence

ChatGPT was used for the final proofread of the manuscript to revise spelling, grammar and style. Alphafold3 and AIUPred were used as detailed above.

## Supporting information

all supplemental materials

## Acknowledgments

We want to gratefully acknowledge Ikuko Hara-Nishimura, Fernando Aniento, Enrique Rojo, Elias Bassil, Marketa Samalova, Karin Schumacher, Rishikesh Bhalerao, Georgia Drakakaki, Michael Sauer, Jiri Friml, Daisaku Ohta for the collegial provision of mutant and transgenic seeds and John Sedbrook for the generous gift of SKU5 antiserum. FUNDING INFORMATION: G.J.S. was supported by the Austrian Science Fund (FWF) grants P 32332-B and P 36778-B. T.A. was supported by Scientific and Technological Research Council of Türkiye (TÜBİTAK) International Postdoctoral Research Fellowship Programme (2024/1), Grant No. 1059B192400105.

## Author contributions

CH, HX and TA conducted research. GJS conceived of the project, conducted research and wrote the paper.

## References

Abramson J, Adler J, Dunger J, Evans R, Green T, Pritzel A, Ronneberger O, Willmore L, Ballard AJ, Bambrick J, Bodenstein SW, Evans DA, Hung CC, O’Neill M, Reiman D, Tunyasuvunakool K, Wu Z, Zemgulyte A, Arvaniti E, Beattie C, Bertolli O, Bridgland A, Cherepanov A, Congreve M, Cowen-Rivers AI, Cowie A, Figurnov M, Fuchs FB, Gladman H, Jain R, Khan YA, Low CMR, Perlin K, Potapenko A, Savy P, Singh S, Stecula A, Thillaisundaram A, Tong C, Yakneen S, Zhong ED, Zielinski M, Zidek A, Bapst V, Kohli P, Jaderberg M, Hassabis D, Jumper JM (2024) Accurate structure prediction of biomolecular interactions with AlphaFold 3. Nature 630 (8016):493–500. doi:10.1038/s41586-024-07487-w

Andeme-Onzighi C, Sivaguru M, Judy-March J, Baskin TI, Driouich A (2002) The *reb1-1* mutation of Arabidopsis alters the morphology of trichoblasts, the expression of arabinogalactan-proteins and the organization of cortical microtubules. Planta 215 (6):949–958

Basu D, Wang W, Ma S, DeBrosse T, Poirier E, Emch K, Soukup E, Tian L, Showalter AM (2015) Two Hydroxyproline Galactosyltransferases, GALT5 and GALT2, Function in Arabinogalactan-Protein Glycosylation, Growth and Development in Arabidopsis. PLoS ONE 10 (5):e0125624. doi:10.1371/journal.pone.0125624

Bernat-Silvestre C, De Sousa Vieira V, Sanchez-Simarro J, Pastor-Cantizano N, Hawes C, Marcote MJ, Aniento F (2020) p24 Family Proteins Are Involved in Transport to the Plasma Membrane of GPI-Anchored Proteins in Plants. Plant Physiol 184 (3):1333–1347. doi:10.1104/pp.20.00880

Borassi C, Gloazzo Dorosz J, Ricardi MM, Carignani Sardoy M, Pol Fachin L, Marzol E, Mangano S, Rodriguez Garcia DR, Martinez Pacheco J, Rondon Guerrero YDC, Velasquez SM, Villavicencio B, Ciancia M, Seifert G, Verli H, Estevez JM (2020) A cell surface arabinogalactan-peptide influences root hair cell fate. New Phytol 227 (3):732–743. doi:10.1111/nph.16487

Borner GH, Lilley KS, Stevens TJ, Dupree P (2003) Identification of glycosylphosphatidylinositol-anchored proteins in arabidopsis. A proteomic and genomic analysis. Plant Physiol 132 (2):568–577

Borner GH, Sherrier DJ, Stevens TJ, Arkin IT, Dupree P (2002) Prediction of glycosylphosphatidylinositol-anchored proteins in arabidopsis. A genomic analysis. Plant Physiol 129 (2):486–499

Boutte Y, Jonsson K, McFarlane HE, Johnson E, Gendre D, Swarup R, Friml J, Samuels L, Robert S, Bhalerao RP (2013) ECHIDNA-mediated post-Golgi trafficking of auxin carriers for differential cell elongation. Proc Natl Acad Sci U S A 110 (40):16259–16264. doi:10.1073/pnas.1309057110

Cunningham BC, Wells JA (1989) High-resolution epitope mapping of hGH-receptor interactions by alanine-scanning mutagenesis. Science 244 (4908):1081–1085. doi:10.1126/science.2471267

Ding L, Zhu JK (1997) A role for arabinogalactan-proteins in root epidermal cell expansion. Planta 203 (3):289–294

Erdos G, Deutsch N, Dosztanyi Z (2025) AIUPred - Binding: Energy Embedding to Identify Disordered Binding Regions. J Mol Biol 437 (15):169071. doi:10.1016/j.jmb.2025.169071

Erdos G, Dosztanyi Z (2024) AIUPred: combining energy estimation with deep learning for the enhanced prediction of protein disorder. Nucleic Acids Res 52 (W1):W176–W181. doi:10.1093/nar/gkae385

Gao H, Zhang Y, Wang W, Zhao K, Liu C, Bai L, Li R, Guo Y (2017) Two Membrane-Anchored Aspartic Proteases Contribute to Pollen and Ovule Development. Plant Physiol 173 (1):219–239. doi:10.1104/pp.16.01719

Gazaway E, Kandel R, Grant OC, Woods RJ (2025) Are N-linked glycans intrinsically disordered? Curr Opin Struct Biol 93:103118. doi:10.1016/j.sbi.2025.103118

Gendre D, McFarlane HE, Johnson E, Mouille G, Sjodin A, Oh J, Levesque-Tremblay G, Watanabe Y, Samuels L, Bhalerao RP (2013) Trans-Golgi network localized ECHIDNA/Ypt interacting protein complex is required for the secretion of cell wall polysaccharides in Arabidopsis. The Plant cell 25 (7):2633–2646. doi:10.1105/tpc.113.112482

Gendre D, Oh J, Boutte Y, Best JG, Samuels L, Nilsson R, Uemura T, Marchant A, Bennett MJ, Grebe M, Bhalerao RP (2011) Conserved Arabidopsis ECHIDNA protein mediates trans-Golgi-network trafficking and cell elongation. P Natl Acad Sci USA 108 (19):8048–8053. doi:10.1073/pnas.1018371108

Goedhart J, von Stetten D, Noirclerc-Savoye M, Lelimousin M, Joosen L, Hink MA, van Weeren L, Gadella TW, Jr., Royant A (2012) Structure-guided evolution of cyan fluorescent proteins towards a quantum yield of 93%. Nat Commun 3:751. doi:10.1038/ncomms1738

Hao C, Zou Q, Bai X, Shi W (2025) Effect of glycosylation on protein folding: From biological roles to chemical protein synthesis. iScience 28 (6):112605. doi:10.1016/j.isci.2025.112605

He Y, Gao J, Luo M, Gao C, Lin Y, Wong HY, Cui Y, Zhuang X, Jiang L (2023) VAMP724 and VAMP726 are involved in autophagosome formation in Arabidopsis thaliana. Autophagy 19 (5):1406–1423. doi:10.1080/15548627.2022.2127240

Heckman JW, Terhune BT, Lamport DT (1988) Characterization of native and modified extensin monomers and oligomers by electron microscopy and gel filtration. Plant Physiol 86 (3):848–856. doi:10.1104/pp.86.3.848

Hijazi M, Durand J, Pichereaux C, Pont F, Jamet E, Albenne C (2012) Characterization of the arabinogalactan protein 31 (AGP31) of *Arabidopsis thaliana*: new advances on the Hyp-*O*-glycosylation of the Pro-rich domain. J Biol Chem 287 (12):9623–9632. doi:10.1074/jbc.M111.247874

Huttner S, Veit C, Schoberer J, Grass J, Strasser R (2012) Unraveling the function of Arabidopsis thaliana OS9 in the endoplasmic reticulum-associated degradation of glycoproteins. Plant molecular biology 79 (1-2):21–33. doi:10.1007/s11103-012-9891-4

Huttner S, Veit C, Vavra U, Schoberer J, Liebminger E, Maresch D, Grass J, Altmann F, Mach L, Strasser R (2014) Arabidopsis Class I alpha-Mannosidases MNS4 and MNS5 Are Involved in Endoplasmic Reticulum-Associated Degradation of Misfolded Glycoproteins. The Plant cell 26 (4):1712–1728. doi:10.1105/tpc.114.123216

Johnson KL, Cassin AM, Lonsdale A, Bacic A, Doblin MS, Schultz CJ (2017a) Pipeline to Identify Hydroxyproline-Rich Glycoproteins. Plant Physiol 174 (2):886–903. doi:10.1104/pp.17.00294

Johnson KL, Cassin AM, Lonsdale A, Wong GK, Soltis DE, Miles NW, Melkonian M, Melkonian B, Deyholos MK, Leebens-Mack J, Rothfels CJ, Stevenson DW, Graham SW, Wang X, Wu S, Pires JC, Edger PP, Carpenter EJ, Bacic A, Doblin MS, Schultz CJ (2017b) Insights into the Evolution of Hydroxyproline-Rich Glycoproteins from 1000 Plant Transcriptomes. Plant Physiol 174 (2):904–921. doi:10.1104/pp.17.00295

Johnson KL, Jones BJ, Bacic A, Schultz CJ (2003) The fasciclin-like arabinogalactan proteins of arabidopsis. A multigene family of putative cell adhesion molecules. Plant Physiol 133 (4):1911–1925

Kaur D, Held MA, Zhang Y, Moreira D, Coimbra S, Showalter AM (2023) Knockout of eight hydroxyproline-O-galactosyltransferases cause multiple vegetative and reproductive growth defects. Cell Surf 10:100117. doi:10.1016/j.tcsw.2023.100117

Kieliszewski MJ (2001) The latest hype on Hyp-*O-*glycosylation codes. Phytochemistry 57 (3):319–323

Kokoszka ME, Kay BK (2015) Mapping protein-protein interactions with phage-displayed combinatorial peptide libraries and alanine scanning. Methods Mol Biol 1248:173–188. doi:10.1007/978-1-4939-2020-4_12

Kurotani A, Tokmakov AA, Kuroda Y, Fukami Y, Shinozaki K, Sakurai T (2014) Correlations between predicted protein disorder and post-translational modifications in plants. Bioinformatics 30 (8):1095–1103. doi:10.1093/bioinformatics/btt762

Lamport DT, Varnai P, Seal CE (2014) Back to the future with the AGP-Ca2+ flux capacitor. Ann Bot 114 (6):1069–1085. doi:10.1093/aob/mcu161

Liu L, Qin L, Safdar LB, Zhao C, Cheng X, Xie M, Zhang Y, Gao F, Bai Z, Huang J, Bhalerao RP, Liu S, Wei Y (2023) The plant trans-Golgi network component ECHIDNA regulates defense, cell death, and endoplasmic reticulum stress. Plant Physiol 191 (1):558–574. doi:10.1093/plphys/kiac400

Lopez-Hernandez F, Tryfona T, Rizza A, Yu X, Harris MO, Webb AA, Kotake T, Dupree P (2020) Calcium Binding by Arabinogalactan Polysaccharides Is Important for Normal Plant Development. Plant Cell. doi:10.1105/tpc.20.00027

Ma Y, Johnson K (2023) Arabinogalactan proteins - Multifunctional glycoproteins of the plant cell wall. Cell Surf 9:100102. doi:10.1016/j.tcsw.2023.100102

Ma Y, MacMillan CP, de Vries L, Mansfield SD, Hao P, Ratcliffe J, Bacic A, Johnson KL (2022) FLA11 and FLA12 glycoproteins fine-tune stem secondary wall properties in response to mechanical stresses. New Phytol 233 (4):1750–1767. doi:10.1111/nph.17898

Ma Y, Shafee T, Mudiyanselage AM, Ratcliffe J, MacMillan CP, Mansfield SD, Bacic A, Johnson KL (2023) Distinct functions of FASCILIN-LIKE ARABINOGALACTAN PROTEINS relate to domain structure. Plant Physiol 192 (1):119–132. doi:10.1093/plphys/kiad097

Miao Y, Wang Y, Grate EVL, Fang X (2026) A Multidimensional View of Biomolecular Condensates in Plant Biology. Annu Rev Plant Biol 77 (1):197–223. doi:10.1146/annurev-arplant-070725-083459

Morrison KL, Weiss GA (2001) Combinatorial alanine-scanning. Curr Opin Chem Biol 5 (3):302–307. doi:10.1016/s1367-5931(00)00206-4

Nguema-Ona E, Andeme-Onzighi C, Aboughe-Angone S, Bardor M, Ishii T, Lerouge P, Driouich A (2006) The *reb1-1* mutation of arabidopsis: Effect on the structure and localization of galactose-containing cell wall polysaccharides. Plant Physiol 140:1406–1417

Ogawa-Ohnishi M, Matsubayashi Y (2015) Identification of three potent hydroxyproline *O-*galactosyltransferases in Arabidopsis. Plant J 81 (5):736–746. doi:10.1111/tpj.12764

Poon S, Heath RL, Clarke AE (2012) A chimeric arabinogalactan protein promotes somatic embryogenesis in cotton cell culture. Plant Physiol 160 (2):684–695. doi:10.1104/pp.112.203075

Qu Y, Egelund J, Gilson PR, Houghton F, Gleeson PA, Schultz CJ, Bacic A (2008) Identification of a novel group of putative *Arabidopsis thaliana* beta-(1,3)-galactosyltransferases. Plant Mol Biol 68 (1-2):43–59. doi:10.1007/s11103-008-9351-3

Rösti J, Barton CJ, Albrecht S, Dupree P, Pauly M, Findlay K, Roberts K, Seifert GJ (2007) UDP-glucose 4-epimerase isoforms UGE2 and UGE4 cooperate in providing UDP-galactose for cell wall biosynthesis and growth of *Arabidopsis thaliana*. Plant Cell 19:1565–1579

Sadeghi H, Birnbaumer M (1999) O-Glycosylation of the V2 vasopressin receptor. Glycobiology 9 (7):731–737. doi:10.1093/glycob/9.7.731

Samalova M, Melnikava A, Elsayad K, Peaucelle A, Gahurova E, Gumulec J, Spyroglou I, Zemlyanskaya EV, Ubogoeva EV, Balkova D, Demko M, Blavet N, Alexiou P, Benes V, Mouille G, Hejatko J (2023) Hormone-regulated expansins: Expression, localization, and cell wall biomechanics in Arabidopsis root growth. Plant Physiol 194 (1):209–228. doi:10.1093/plphys/kiad228

Satpute-Krishnan P, Ajinkya M, Bhat S, Itakura E, Hegde RS, Lippincott-Schwartz J (2014) ER stress-induced clearance of misfolded GPI-anchored proteins via the secretory pathway. Cell 158 (3):522–533. doi:10.1016/j.cell.2014.06.026

Sedbrook JC, Carroll KL, Hung KF, Masson PH, Somerville CR (2002) The Arabidopsis SKU5 gene encodes an extracellular glycosyl phosphatidylinositol-anchored glycoprotein involved in directional root growth. Plant Cell 14 (7):1635–1648

Seifert GJ (2020) On the Potential Function of Type II Arabinogalactan O-Glycosylation in Regulating the Fate of Plant Secretory Proteins. Front Plant Sci 11:563735. doi:10.3389/fpls.2020.563735

Seifert GJ, Barber C, Wells B, Dolan L, Roberts K (2002) Galactose biosynthesis in arabidopsis: genetic evidence for substrate channeling from UDP-D-galactose into cell wall polymers. Curr Biol 12 (21):1840–1845

Shafee T, Bacic A, Johnson K (2020) Evolution of sequence-diverse disordered regions in a protein family: order within the chaos. Mol Biol Evol. doi:10.1093/molbev/msaa096

Showalter AM, Basu D (2016) Glycosylation of arabinogalactan-proteins essential for development in Arabidopsis. Commun Integr Biol 9 (3):e1177687

Showalter AM, Keppler B, Lichtenberg J, Gu D, Welch LR (2010) A bioinformatics approach to the identification, classification, and analysis of hydroxyproline-rich glycoproteins. Plant Physiol 153 (2):485–513. doi:10.1104/pp.110.156554

Shpak E, Barbar E, Leykam JF, Kieliszewski MJ (2001) Contiguous hydroxyproline residues direct hydroxyproline arabinosylation in *Nicotiana tabacum*. J Biol Chem 276 (14):11272–11278

Sikorska N, Lemus L, Aguilera-Romero A, Manzano-Lopez J, Riezman H, Muniz M, Goder V (2016) Limited ER quality control for GPI-anchored proteins. The Journal of cell biology 213 (6):693–704. doi:10.1083/jcb.201602010

Spurway TD, Dalley JA, High S, Bulleid NJ (2001) Early events in glycosylphosphatidylinositol anchor addition. substrate proteins associate with the transamidase subunit gpi8p. J Biol Chem 276 (19):15975–15982. doi:10.1074/jbc.M010128200

Stafstrom JP, Staehelin LA (1986) The role of carbohydrate in maintaining extensin in an extended conformation. Plant Physiol 81 (1):242–246. doi:10.1104/pp.81.1.242

Stoddard A, Rolland V (2019) I see the light! Fluorescent proteins suitable for cell wall/apoplast targeting in Nicotiana benthamiana leaves. Plant Direct 3 (1):e00112. doi:10.1002/pld3.112

Strasser R (2016) Plant protein glycosylation. Glycobiology 26 (9):926–939. doi:10.1093/glycob/cww023

Strasser R (2022) Recent Developments in Deciphering the Biological Role of Plant Complex N-Glycans. Front Plant Sci 13:897549. doi:10.3389/fpls.2022.897549

Strasser R, Seifert G, Doblin MS, Johnson KL, Ruprecht C, Pfrengle F, Bacic A, Estevez JM (2021) Cracking the “Sugar Code”: A Snapshot of N- and O-Glycosylation Pathways and Functions in Plants Cells. Front Plant Sci 12:640919. doi:10.3389/fpls.2021.640919

Tan L, Eberhard S, Pattathil S, Warder C, Glushka J, Yuan C, Hao Z, Zhu X, Avci U, Miller JS, Baldwin D, Pham C, Orlando R, Darvill A, Hahn MG, Kieliszewski MJ, Mohnen D (2013) An Arabidopsis cell wall proteoglycan consists of pectin and arabinoxylan covalently linked to an arabinogalactan protein. Plant Cell 25 (1):270–287. doi:10.1105/tpc.112.107334

Tan L, Leykam JF, Kieliszewski MJ (2003) Glycosylation motifs that direct arabinogalactan addition to arabinogalactan-proteins. Plant Physiol 132 (3):1362–1369

Tan L, Zhang L, Black I, Glushka J, Urbanowicz B, Heiss C, Azadi P (2023) Most of the rhamnogalacturonan-I from cultured Arabidopsis cell walls is covalently linked to arabinogalactan-protein. Carbohydr Polym 301 (Pt B):120340. doi:10.1016/j.carbpol.2022.120340

Teh OK, Singh P, Ren J, Huang LT, Ariyarathne M, Salamon BP, Wang Y, Kotake T, Fujita T (2022) Surface-localized glycoproteins act through class C ARFs to fine-tune gametophore initiation in Physcomitrium patens. Development 149 (24). doi:10.1242/dev.200370

Theillet FX, Kalmar L, Tompa P, Han KH, Selenko P, Dunker AK, Daughdrill GW, Uversky VN (2013) The alphabet of intrinsic disorder: I. Act like a Pro: On the abundance and roles of proline residues in intrinsically disordered proteins. Intrinsically Disord Proteins 1 (1):e24360. doi:10.4161/idp.24360

Uversky VN (2017) Paradoxes and wonders of intrinsic disorder: Stability of instability. Intrinsically Disord Proteins 5 (1):e1327757. doi:10.1080/21690707.2017.1327757

van der Hoorn RA (2008) Plant proteases: from phenotypes to molecular mechanisms. Annu Rev Plant Biol 59:191–223. doi:10.1146/annurev.arplant.59.032607.092835

Velasquez SM, Marzol E, Borassi C, Pol-Fachin L, Ricardi MM, Mangano S, Juarez SP, Salter JD, Dorosz JG, Marcus SE, Knox JP, Dinneny JR, Iusem ND, Verli H, Estevez JM (2015) Low Sugar Is Not Always Good: Impact of Specific O-Glycan Defects on Tip Growth in Arabidopsis. Plant Physiology 168 (3):808–813. doi:10.1104/pp.114.255521

Williams RM, Obradovi Z, Mathura V, Braun W, Garner EC, Young J, Takayama S, Brown CJ, Dunker AK (2001) The protein non-folding problem: amino acid determinants of intrinsic order and disorder. Pac Symp Biocomput:89–100. doi:10.1142/9789814447362_0010

Xiao L, Xia K (2025) Functions of Intrinsically Disordered Regions. Biology (Basel) 14 (7). doi:10.3390/biology14070810

Xu SL, Rahman A, Baskin TI, Kieber JJ (2008) Two leucine-rich repeat receptor kinases mediate signaling, linking cell wall biosynthesis and ACC synthase in Arabidopsis. Plant Cell 20 (11):3065–3079

Xue H, Veit C, Abas L, Tryfona T, Maresch D, Ricardi MM, Estevez JM, Strasser R, Seifert GJ (2017) Arabidopsis thaliana FLA4 functions as a glycan-stabilized soluble factor via its carboxy-proximal Fasciclin 1 domain. Plant J 91 (4):613–630. doi:10.1111/tpj.13591

Zhang N, Wright T, Wang X, Karki U, Savary BJ, Xu J (2019) Engineering ‘designer’ glycomodules for boosting recombinant protein secretion in tobacco hairy root culture and studying hydroxyproline-*O*-glycosylation process in plants. Plant Biotechnol J 17 (6):1130–1141. doi:10.1111/pbi.13043

Zurzolo C, Simons K (2016) Glycosylphosphatidylinositol-anchored proteins: Membrane organization and transport. Biochimica et biophysica acta 1858 (4):632–639. doi:10.1016/j.bbamem.2015.12.018

