## Supplementary material for "An intrinsically disordered proline-rich domain acts in synergy with AGP-type glycosylation to control the protein fate of FLA4": all supplemental materials: all supplemental materials.docx

**supplemental Table S1.** FLA4 constructs used in this study. only the Pro-rich domains are shown in detail.

```
WT	     NSP.Fas1-1.LTPPPTSTSLSPPPAG.Fas1-2.LPKELFPKSGQPVATAPPQEISLSPESSSEQPSRLVSPPREIVS.GPI-signal
P1234A     NSP.Fas1-1.LTaaaTSTSLSaaaAG.Fas1-2.LPKELFPKSGQPVATAaaQEISLSPESSSEQPSRLVSaaREIVS.GPI-signal
P12A	     NSP.Fas1-1.LTaaaTSTSLSaaaAG.Fas1-2.LPKELFPKSGQPVATAPPQEISLSPESSSEQPSRLVSPPREIVS.GPI-signal
P34A	     NSP.Fas1-1.LTPPPTSTSLSPPPAG.Fas1-2.LPKELFPKSGQPVATAaaQEISLSPESSSEQPSRLVSaaREIVS.GPI-signal
P8A        NSP.Fas1-1.LTPPPTSTSLSPPPAG.Fas1-2.LPKELFaKSGQaVATAaaQEISLSaESSSEQaSRLVSaaREIVS.GPI-signal
P3A	     NSP.Fas1-1.LTPPPTSTSLSPPPAG.Fas1-2.LPKELFPKSGQPVATAaaQEISLSPESSSEQPSRLVSPPREIVS.GPI-signal
P4A	     NSP.Fas1-1.LTPPPTSTSLSPPPAG.Fas1-2.LPKELFPKSGQPVATAPPQEISLSPESSSEQPSRLVSaaREIVS.GPI-signal
P6A        NSP.Fas1-1.LTPPPTSTSLSPPPAG.Fas1-2.LPKELFaKSGQaVATAaaQEISLSaESSSEQaSRLVSPPREIVS.GPI-signal
PR2_AGmod  NSP.Fas1-1.LTPPPTSTSLSPPPAG.Fas1-2.LPKELFPKSGQPVATAPPQEISLSPspspEQPSRLVSPPREIVS.GPI-signal
P8A_AGmod  NSP.Fas1-1.LTPPPTSTSLSPPPAG.Fas1-2.LPKELFaKSGQaVATAaaQEISLSPspspEQaSRLVSaaREIVS.GPI-signal
P8A_3xNGly NSP.Fas1-1.LTPPPTSTSLSPPPAG.Fas1-2.LPKELFnKSGQaVATAaaQEISLSnESSSEQaSvngSaaREIVS.GPI-signal
PR2toEXT   NSP.Fas1-1.LTPPPTSTSLSPPPAG.Fas1-2.LPKHYSPPPVYKSPPPPVKHYSPPPVYKSPPPPVKHYSPPPVYKSGPI-signal
P34G	     NSP.Fas1-1.LTPPPTSTSLSPPPAG.Fas1-2.LPKELFPKSGQPVATAggQEISLSPESSSEQPSRLVSggREIVS.GPI-signal
PR2_10xG    all as above     Fas1-2.LPKELFPKSGQPVATAPPQEISLSPESSGGGGGGGGGGSEQPSRLVSPPREIVSGPI-signal
```

**supplemental table S2: genetic analysis of F4T_P8A fate**

| **Mutant locus** | **cellular role of protein** | **references** | **confocal**  **observation of F4T_P8A or F4C_P8A** | **Western blot ±CHX** |
| --- | --- | --- | --- | --- |
| **ER to Golgi, ERAD** | | | |  |
| *mag2 mip3* | Golgi-ER retrograde transport | (Zhao et al. 2018; Li et al. 2013) | partial ER retention | n.e. |
| *p24δ3δ4δ5δ6* | GPI-AP ER-export | (Bernat-Silvestre et al. 2020) | ER-retention | reduced degradation |
| *mns4 mns5* | ERAD | (Huttner et al. 2014) | n.e. | n.e. |
| **TGN function and cargo/receptor/tethering complexes** | | | |  |
| *mtv16-1/vps51-1* | GARP^†)^ | (Delgadillo et al. 2020) | increased signal in apoplast | n.t. ^‡^ |
| *nhx5 nhx6* | TGN&PVC K^+^:H^+^ antiporter | (Reguera et al. 2015) | n.e. | n.e. |
| *det3* | TGN H^+^-ATPase | (Brux et al. 2008) | n.e.^§)^ | n.e. |
| *echidna* | TGN/EE function | (Gendre et al. 2013) | partial re-direction to internal compartment | stabilizing |
| *rog2-2, rog2-7* | TGN/EE function | (Kohorn et al. 2016; Rosquete et al. 2019) | n.e. | n.e. |
| *epsin mtv1* | TGN/EE function | (Heinze et al. 2020) | *Increased* vacuolar targeting (both F4T and F4T_P8A) | n.t. ^‡^ |
| *trs120-4* | TRAPPII^*)^ | (Thellmann et al. 2010; Qi et al. 2011) | n.e. | n.t. ^‡^ |
| *vsr5 vsr6 vsr7* | cargo receptors | (Lee et al. 2013) | n.e. | n.e. |
| **Routing to vacuolar compartment** | | | |  |
| *mtv14-3/gfs10-2* | unknown | (Delgadillo et al. 2020) | n.e. | n.e. |
| *atg5-1* | core autophagy |  | n.e. | n.e. |
| *mtv11-1/vps15-1* | PI3K subunit | (Delgadillo et al. 2020) | n.e. | n.e. |
| *pten2a pten2b* | PI3Pase | (Pribat et al. 2012; Delgadillo et al. 2020) | n.e. | n.e. |
| *vps9-2* | RAB5 GEF | (Goh et al. 2007) | n.e. | n.e. |
| *mtv7-2/erel1-1* | RAB5-effector | (Sakurai et al. 2016; Delgadillo et al. 2020) | n.e. | n.e. |
| *mtv9-2* | RAB5-effector | (Delgadillo et al. 2020; Ito et al. 2018) | Increased apoplastic signal | n.e. |
| *snx1-2* | retromer | (Zhang et al. 2022; Kleine-Vehn et al. 2008) | n.e. | n.e. |
| *vps29-3* | retromer | (Yuan et al. 2018; Kleine-Vehn et al. 2008; Jaillais et al. 2007) | n.e. | n.t. ^‡^ |
| **Endocytosis to vacuole** | | | |  |
| *Auxilin1/2-OE* | endocytosis | (Adamowski et al. 2018) | Reduction of signal | n.t. ^‡^ |
| *alix-1* | ESCRT & retromer | (Bassham 2022) | Increased apoplastic signal | n.t. ^‡^ |
| **Lipids and microdomains** | | | |  |
| *gpi8-1* | GPI-anchor attachment | (Bundy et al. 2016) | n.e. | n.t. ^‡^ |
| *smt2 smt3* | C24-ethylsterol biosynthesis | (Nakamoto et al. 2015) | weaker signal | n.t. ^‡^ |
| *trn2/tet1* | unknown/lipid raft associated | (Head et al. 2014; Wang et al. 2012) | partial re-direction to internal compartment | n.t. ^‡^ |

n.e. : no effect on F4T_P8A compared to the control (*sos5-2* background)

n.t. : not tested

^†^) tethering in TGN to vacuole transport

^‡^) only microscopic analyses were performed for the indicated genotypes (mostly due to infertility/lethality of some mutants or otherwise insufficient seed stocks of homozygous lines))

^§^) the *det3* background caused increased fluorescence of reporters with citrin but not with mTurq tagged FLA4, in both PM/apoplastic and intracellular fraction indicates that endosomal and apoplastic pH might be relatively higher in *det3* compared to wild type.

**References for Table 2**

Adamowski M, Narasimhan M, Kania U, Glanc M, De Jaeger G, Friml J (2018) A Functional Study of AUXILIN-LIKE1 and 2, Two Putative Clathrin Uncoating Factors in Arabidopsis. Plant Cell 30 (3):700-716. doi:10.1105/tpc.17.00785

Bassham DC (2022) An unexpected function for an ESCRT protein. Proc Natl Acad Sci U S A 119 (26):e2207055119. doi:10.1073/pnas.2207055119

Bernat-Silvestre C, De Sousa Vieira V, Sanchez-Simarro J, Pastor-Cantizano N, Hawes C, Marcote MJ, Aniento F (2020) p24 Family Proteins Are Involved in Transport to the Plasma Membrane of GPI-Anchored Proteins in Plants. Plant Physiol 184 (3):1333-1347. doi:10.1104/pp.20.00880

Brux A, Liu TY, Krebs M, Stierhof YD, Lohmann JU, Miersch O, Wasternack C, Schumacher K (2008) Reduced V-ATPase activity in the trans-Golgi network causes oxylipin-dependent hypocotyl growth Inhibition in Arabidopsis. Plant Cell 20 (4):1088-1100. doi:10.1105/tpc.108.058362

Bundy MG, Kosentka PZ, Willet AH, Zhang L, Miller E, Shpak ED (2016) A Mutation in the Catalytic Subunit of the Glycosylphosphatidylinositol Transamidase Disrupts Growth, Fertility, and Stomata Formation. Plant Physiology 171 (2):974-985. doi:10.1104/pp.16.00339

Delgadillo MO, Ruano G, Zouhar J, Sauer M, Shen J, Lazarova A, Sanmartin M, Lai LTF, Deng C, Wang P, Hussey PJ, Sanchez-Serrano JJ, Jiang L, Rojo E (2020) MTV proteins unveil ER- and microtubule-associated compartments in the plant vacuolar trafficking pathway. Proc Natl Acad Sci U S A 117 (18):9884-9895. doi:10.1073/pnas.1919820117

Gendre D, McFarlane HE, Johnson E, Mouille G, Sjodin A, Oh J, Levesque-Tremblay G, Watanabe Y, Samuels L, Bhalerao RP (2013) Trans-Golgi network localized ECHIDNA/Ypt interacting protein complex is required for the secretion of cell wall polysaccharides in Arabidopsis. The Plant cell 25 (7):2633-2646. doi:10.1105/tpc.113.112482

Goh T, Uchida W, Arakawa S, Ito E, Dainobu T, Ebine K, Takeuchi M, Sato K, Ueda T, Nakano A (2007) VPS9a, the common activator for two distinct types of Rab5 GTPases, is essential for the development of Arabidopsis thaliana. Plant Cell 19 (11):3504-3515. doi:10.1105/tpc.107.053876

Head BP, Patel HH, Insel PA (2014) Interaction of membrane/lipid rafts with the cytoskeleton: impact on signaling and function: membrane/lipid rafts, mediators of cytoskeletal arrangement and cell signaling. Biochim Biophys Acta 1838 (2):532-545. doi:10.1016/j.bbamem.2013.07.018

Heinze L, Freimuth N, Rossling AK, Hahnke R, Riebschlager S, Frohlich A, Sampathkumar A, McFarlane HE, Sauer M (2020) EPSIN1 and MTV1 define functionally overlapping but molecularly distinct trans-Golgi network subdomains in Arabidopsis. Proc Natl Acad Sci U S A 117 (41):25880-25889. doi:10.1073/pnas.2004822117

Huttner S, Veit C, Vavra U, Schoberer J, Liebminger E, Maresch D, Grass J, Altmann F, Mach L, Strasser R (2014) Arabidopsis Class I alpha-Mannosidases MNS4 and MNS5 Are Involved in Endoplasmic Reticulum-Associated Degradation of Misfolded Glycoproteins. The Plant cell 26 (4):1712-1728. doi:10.1105/tpc.114.123216

Ito E, Ebine K, Choi SW, Ichinose S, Uemura T, Nakano A, Ueda T (2018) Integration of two RAB5 groups during endosomal transport in plants. Elife 7. doi:10.7554/eLife.34064

Jaillais Y, Santambrogio M, Rozier F, Fobis-Loisy I, Miege C, Gaude T (2007) The retromer protein VPS29 links cell polarity and organ initiation in plants. Cell 130 (6):1057-1070. doi:10.1016/j.cell.2007.08.040

Kleine-Vehn J, Leitner J, Zwiewka M, Sauer M, Abas L, Luschnig C, Friml J (2008) Differential degradation of PIN2 auxin efflux carrier by retromer-dependent vacuolar targeting. P Natl Acad Sci USA 105 (46):17812-17817. doi:10.1073/pnas.0808073105

Kohorn BD, Hoon D, Minkoff BB, Sussman MR, Kohorn SL (2016) Rapid Oligo-Galacturonide Induced Changes in Protein Phosphorylation in Arabidopsis. Mol Cell Proteomics 15 (4):1351-1359. doi:10.1074/mcp.M115.055368

Lee Y, Jang M, Song K, Kang H, Lee MH, Lee DW, Zouhar J, Rojo E, Sohn EJ, Hwang I (2013) Functional identification of sorting receptors involved in trafficking of soluble lytic vacuolar proteins in vegetative cells of Arabidopsis. Plant Physiol 161 (1):121-133. doi:10.1104/pp.112.210914

Li L, Shimada T, Takahashi H, Koumoto Y, Shirakawa M, Takagi J, Zhao X, Tu B, Jin H, Shen Z, Han B, Jia M, Kondo M, Nishimura M, Hara-Nishimura I (2013) MAG2 and three MAG2-INTERACTING PROTEINs form an ER-localized complex to facilitate storage protein transport in Arabidopsis thaliana. Plant J 76 (5):781-791. doi:10.1111/tpj.12347

Nakamoto M, Schmit AC, Heintz D, Schaller H, Ohta D (2015) Diversification of sterol methyltransferase enzymes in plants and a role for beta-sitosterol in oriented cell plate formation and polarized growth. Plant J 84 (5):860-874. doi:10.1111/tpj.13043

Pribat A, Sormani R, Rousseau-Gueutin M, Julkowska MM, Testerink C, Joubes J, Castroviejo M, Laguerre M, Meyer C, Germain V, Rothan C (2012) A novel class of PTEN protein in Arabidopsis displays unusual phosphoinositide phosphatase activity and efficiently binds phosphatidic acid. Biochem J 441 (1):161-171. doi:10.1042/BJ20110776

Qi X, Kaneda M, Chen J, Geitmann A, Zheng H (2011) A specific role for Arabidopsis TRAPPII in post-Golgi trafficking that is crucial for cytokinesis and cell polarity. Plant J 68 (2):234-248. doi:10.1111/j.1365-313X.2011.04681.x

Reguera M, Bassil E, Tajima H, Wimmer M, Chanoca A, Otegui MS, Paris N, Blumwald E (2015) pH Regulation by NHX-Type Antiporters Is Required for Receptor-Mediated Protein Trafficking to the Vacuole in Arabidopsis. Plant Cell 27 (4):1200-1217. doi:10.1105/tpc.114.135699

Rosquete MR, Worden N, Ren G, Sinclair RM, Pfleger S, Salemi M, Phinney BS, Domozych D, Wilkop T, Drakakaki G (2019) AtTRAPPC11/ROG2: A Role for TRAPPs in Maintenance of the Plant Trans-Golgi Network/Early Endosome Organization and Function. Plant Cell 31 (8):1879-1898. doi:10.1105/tpc.19.00110

Sakurai HT, Inoue T, Nakano A, Ueda T (2016) ENDOSOMAL RAB EFFECTOR WITH PX-DOMAIN, an Interacting Partner of RAB5 GTPases, Regulates Membrane Trafficking to Protein Storage Vacuoles in Arabidopsis. Plant Cell 28 (6):1490-1503. doi:10.1105/tpc.16.00326

Thellmann M, Rybak K, Thiele K, Wanner G, Assaad FF (2010) Tethering factors required for cytokinesis in Arabidopsis. Plant Physiol 154 (2):720-732. doi:10.1104/pp.110.154286

Wang F, Vandepoele K, Van Lijsebettens M (2012) Tetraspanin genes in plants. Plant Sci 190:9-15. doi:10.1016/j.plantsci.2012.03.005

Yuan R, Lan J, Fang Y, Yu H, Zhang J, Huang J, Qin G (2018) The Arabidopsis USL1 controls multiple aspects of development by affecting late endosome morphology. New Phytol 219 (4):1388-1405. doi:10.1111/nph.15249

Zhang Y, Wang LF, Han SY, Ren F, Liu WC (2022) Sorting Nexin1 negatively modulates phosphate uptake by facilitating Phosphate Transporter1;1 degradation in Arabidopsis. Plant J 111 (1):72-84. doi:10.1111/tpj.15778

Zhao X, Guo X, Tang X, Zhang H, Wang M, Kong Y, Zhang X, Zhao Z, Lv M, Li L (2018) Misregulation of ER-Golgi Vesicle Transport Induces ER Stress and Affects Seed Vigor and Stress Response. Front Plant Sci 9:658. doi:10.3389/fpls.2018.00658

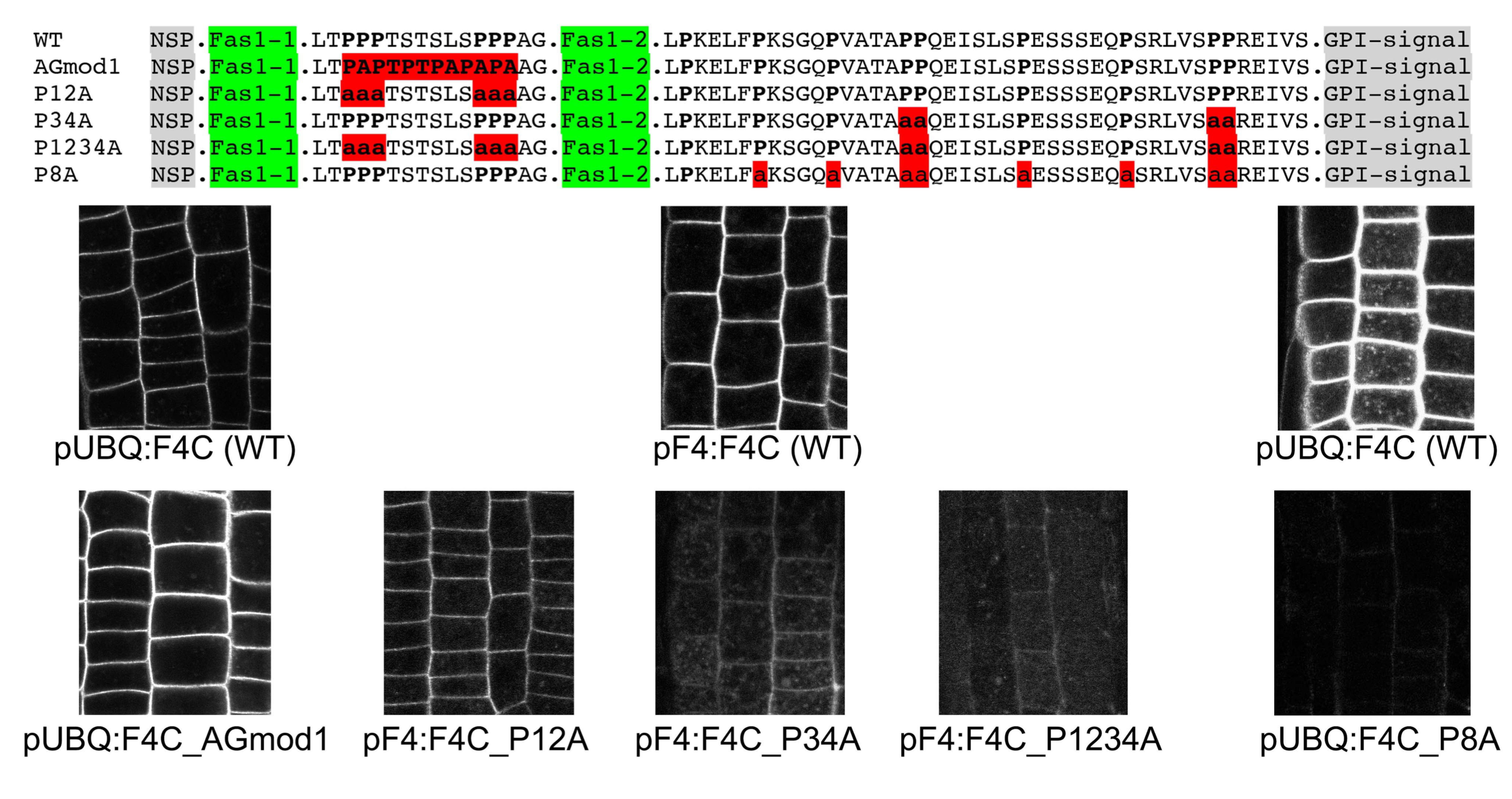

**supplemental Figure S1:** Pro-to-Ala substitutions in the central and the C-terminal Pro-rich domain have different effects on protein localization.

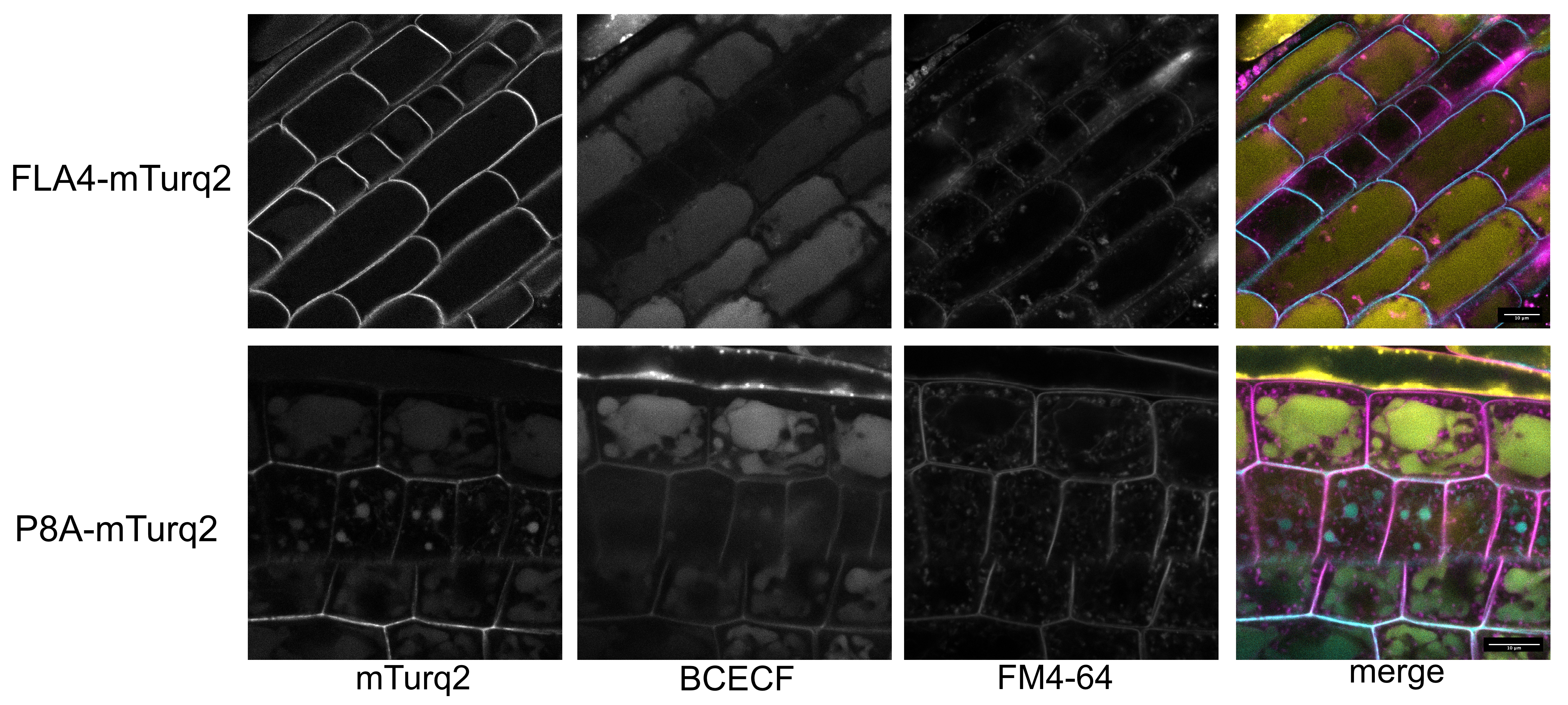

**supplemental Figure S2:** Co-localization of FLA4 reporter constructs with plasma membrane and vacuolar marker dyes

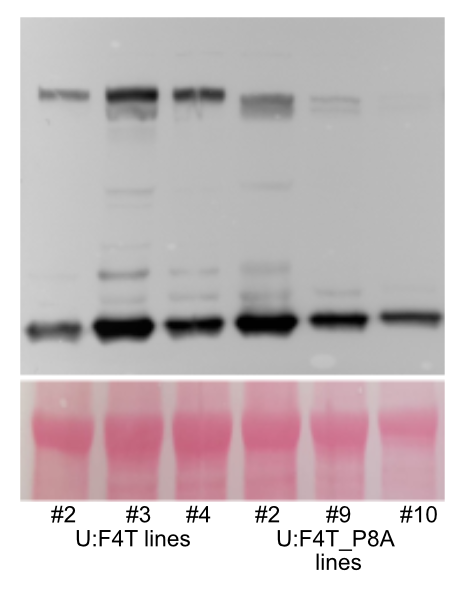

**supplemental Figure S3:** Steady-state protein abundance of independent transformant lines of 'wild type' F4T and Ala-to-Pro substitution variant F4T_P8A. Note that the release of the citrin-tag from the fusion protein is the consequence of pre-electrophoresis heat treatment necessary for SDS-PAGE.

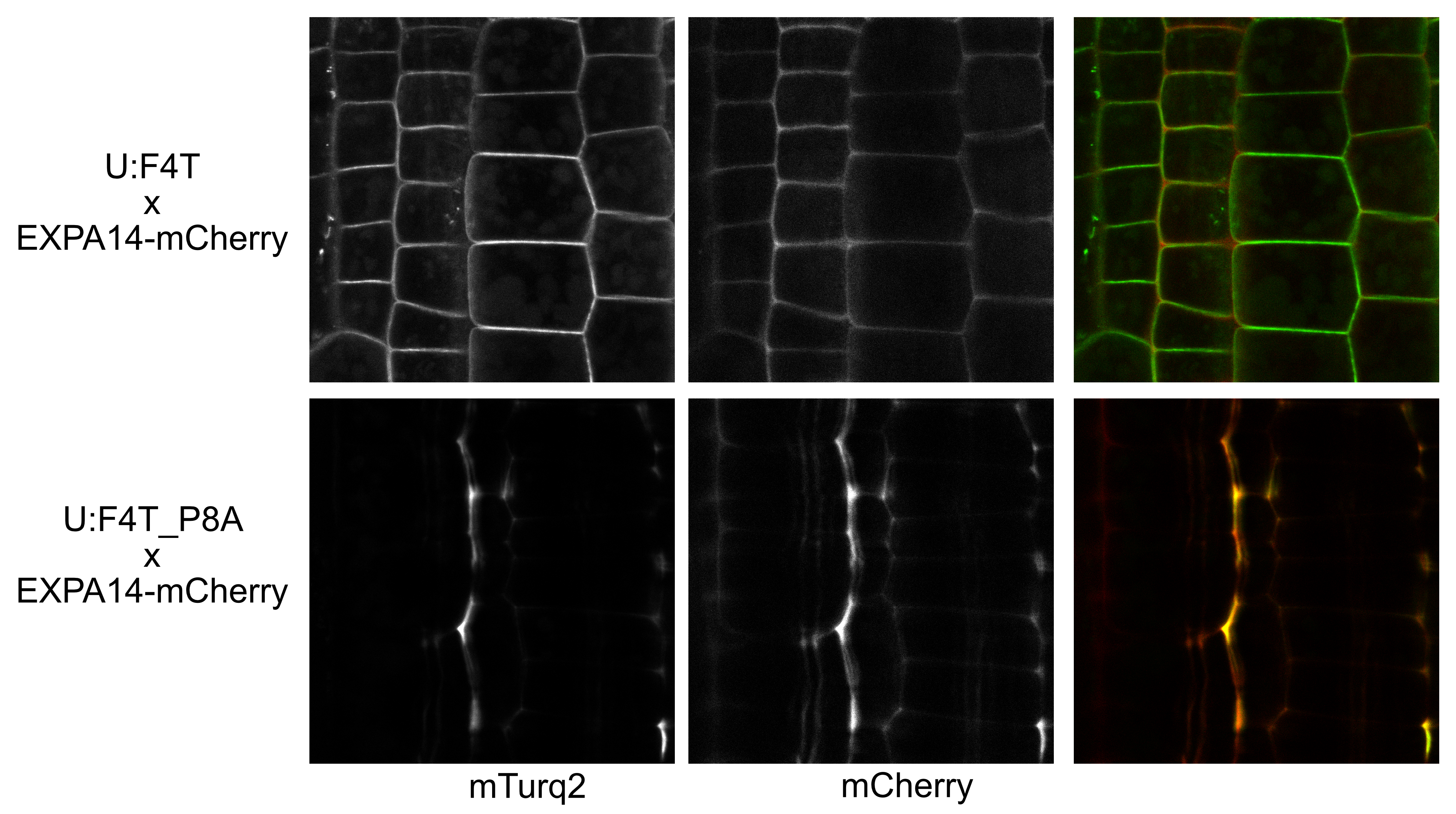
**supplemental Figure S4:** Co-localization of F4T_P8A with the apoplastic reporter EXPA14-mCherry. Note that EXPA14-mCherry can give the impression of a plasma membrane localization in positions of very close proximity between cells but that as opposed to a plasma membrane protein such as F4T, it is localized in cell corners (three-way junctions).

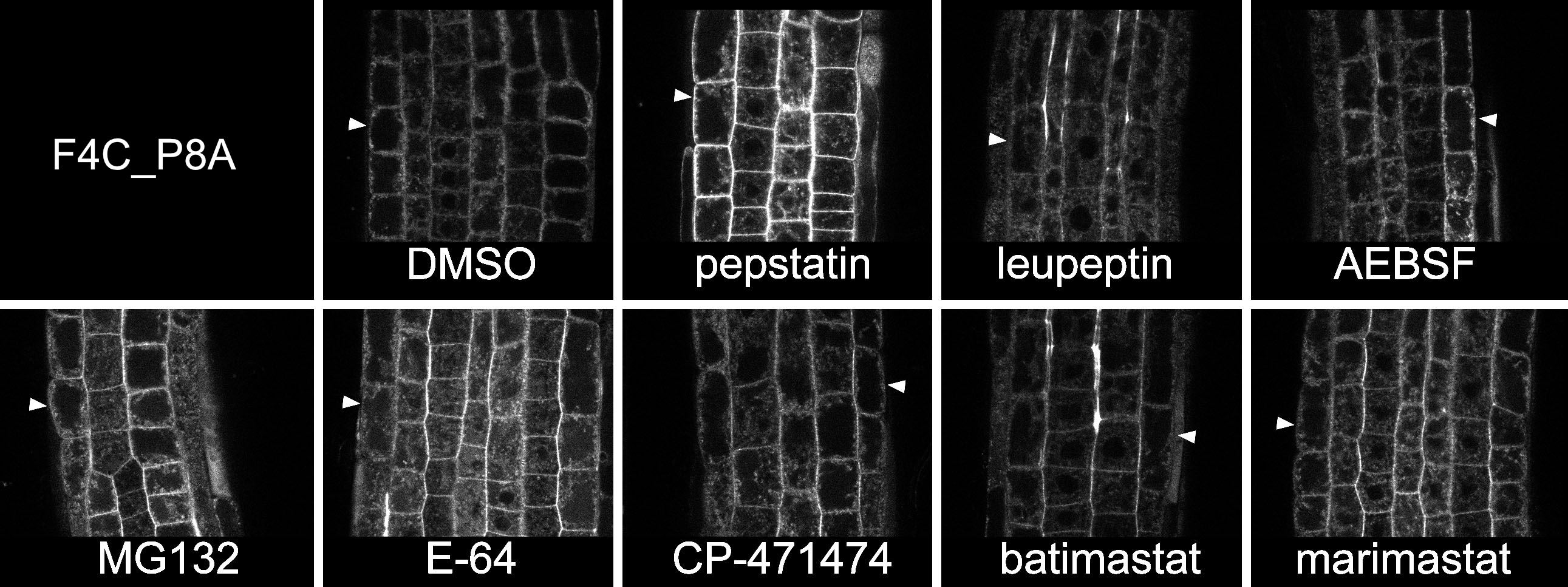

**supplemental Figure S5:** Chemical screen for protease inhibitors affecting the localization of F4C_P8A. Note that only after pepstatinA treatment (10µM 16 h), the reporter is seen at the epidermal plasma membrane (arrowheads).

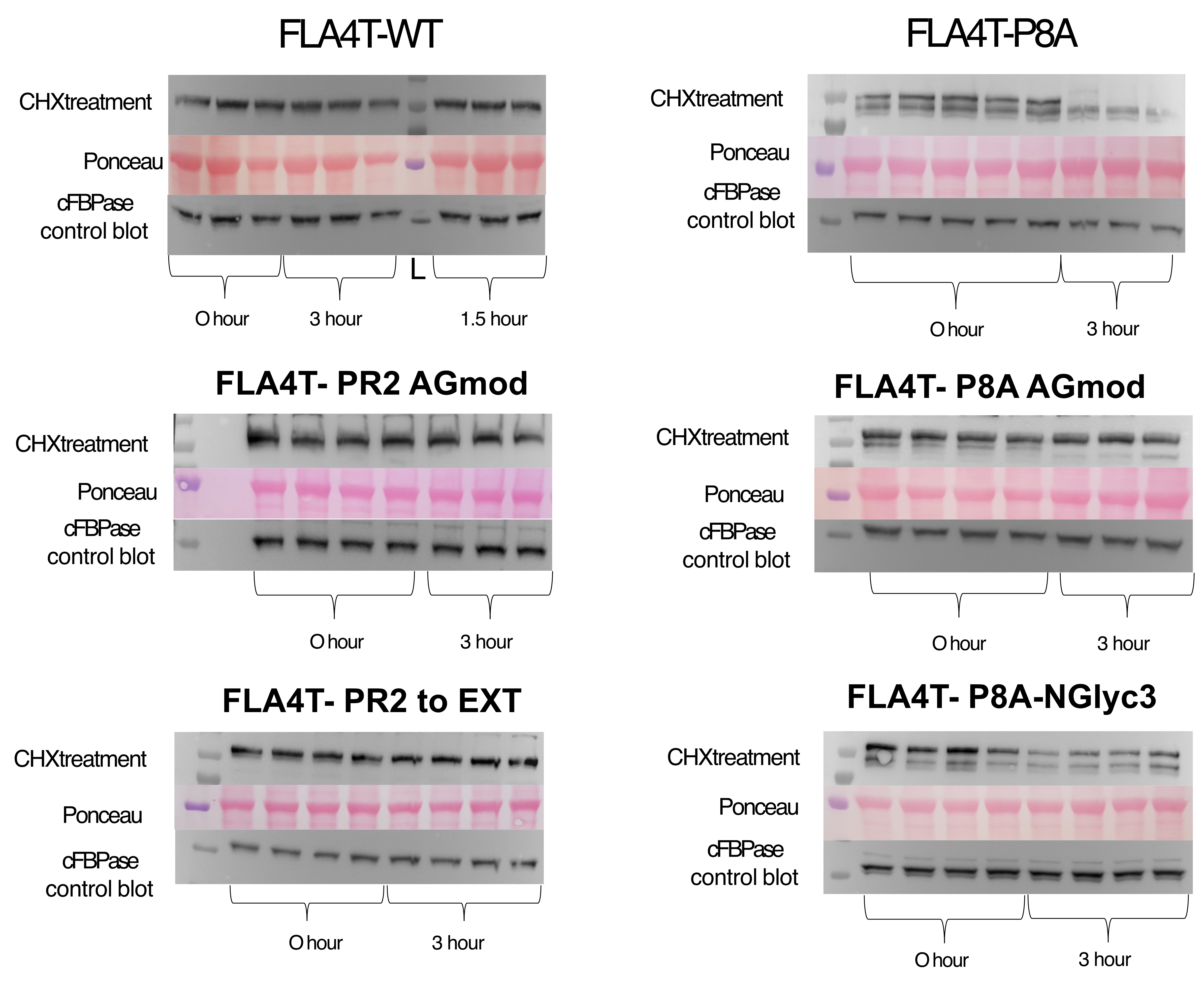

**supplemental Figure S6:** *In planta* stability test for various FLA5 substitution variants*.* In this experiment several technical replicates (same transformant, independent treeatments) are shown.

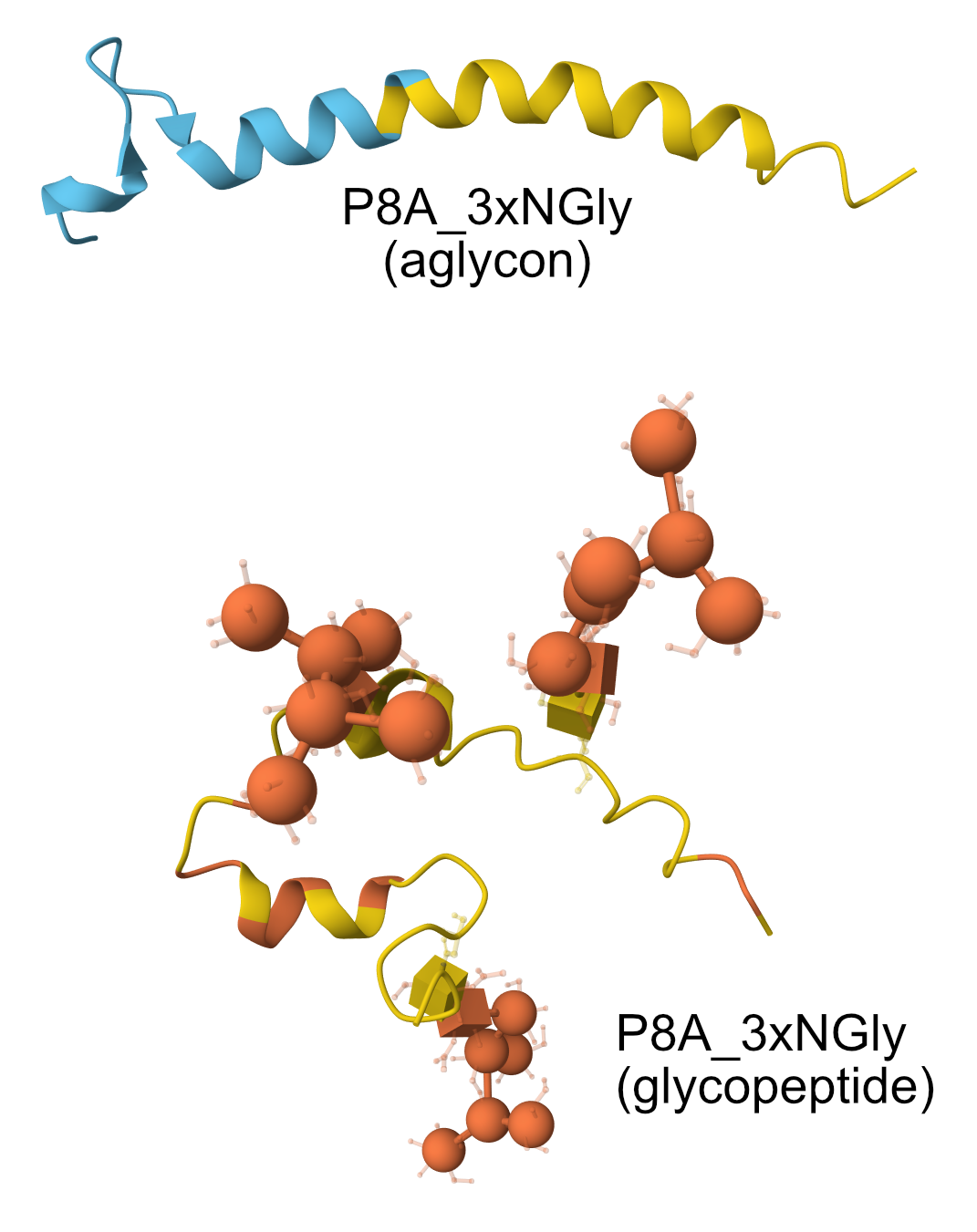

**supplemental Figure S7:** Alphafold3 models of P8A_3xNGly modeled without and including *N-*glycan. Presence of the glycan is predicted to interfere with α-helix formation.

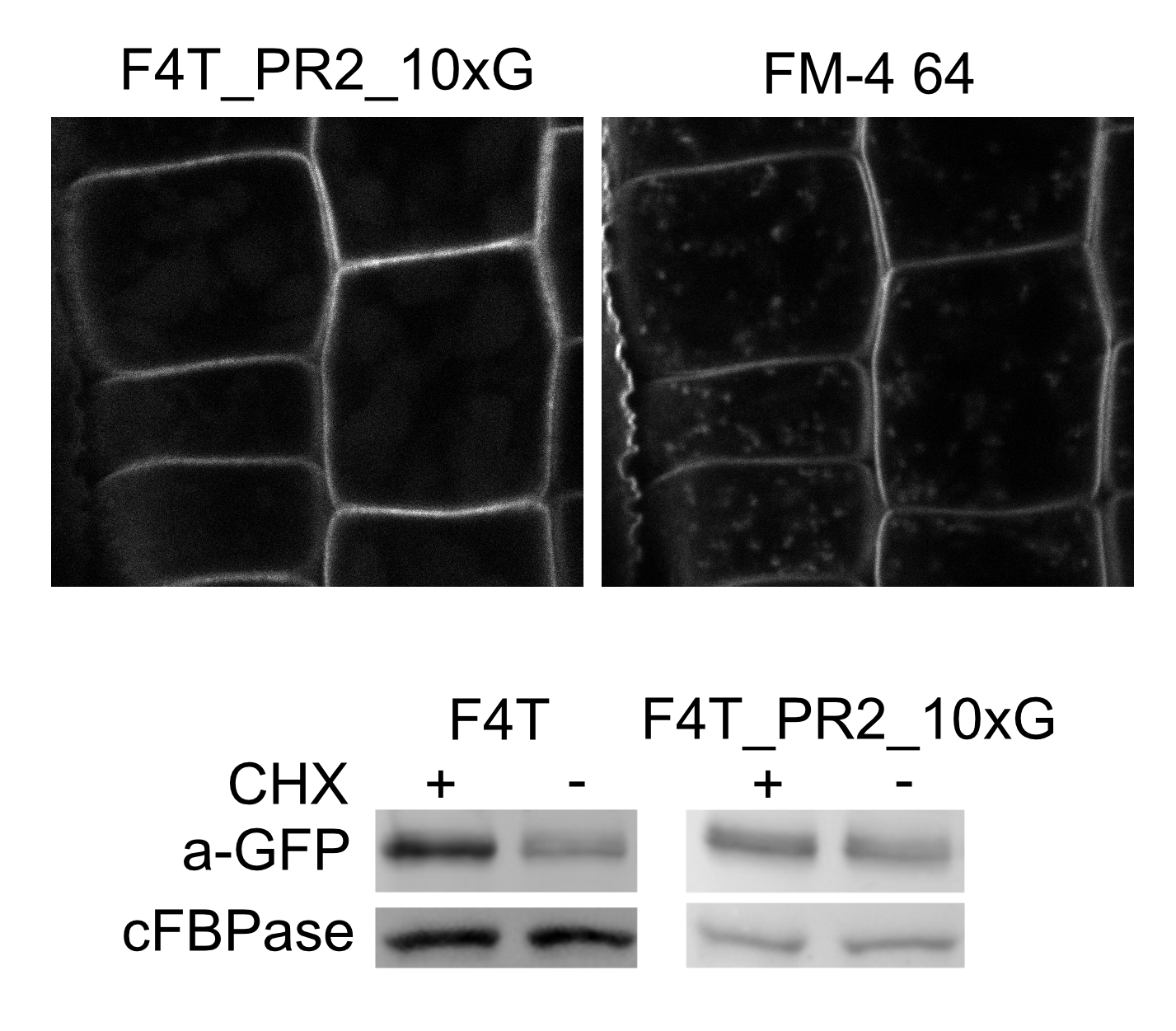

**supplemental Figure S8:** Introduction of a 10 x Gly linker has no negative effect on protein localization or stability.
